# High fat diet but not glucocorticoid-induced obesity results in fatty liver

**DOI:** 10.1101/2025.04.03.647107

**Authors:** Agnieszka Agas, Sanjeev Sharma, Anthony Narciso, Jijo Wilson, Erin Kilkenny, Louise Lantier, Ahsan Uddin, Roma Patel, Timothy E. McGraw, Mary N. Teruel

**Author notes:** Equal contribution (Sanjeev Sharma, Anthony Narciso). Correspondence and requests for materials should be addressed to M.N.T.

## Abstract

Chronic stress, modeled by flattened circadian glucocorticoid (GC) oscillations, is increasingly implicated in obesity. We investigated the underlying mechanisms by disrupting GC rhythms in mice, elevating normally low GC levels during the rest period. This disruption induced substantial obesity, comparable to a 60% high-fat diet (HFD), with additive effects on fat mass, suggesting distinct mechanisms driving adiposity. Despite similar adiposity, GC-flattening and HFD produced profoundly different metabolic outcomes. HFD led to hepatic steatosis and elevated fasting glucose/fatty acid levels, reflecting typical diet-induced dysfunction. In contrast, GC-flattening maintained low fasting glucose/fatty acid levels and prevented hepatic lipid accumulation, with increased adiposity driven by a shift of glucose uptake from muscle to fat and suppression of lipolysis. This reveals a previously unrecognized mechanism of obesity development where excess fat accumulation occurs independently of the mechanisms driving metabolic dysfunction observed in diet-induced obesity. The dissociation between obesity and metabolic dysfunction in GC-flattening challenges the view that increased adiposity and persistent hyperinsulinemia inevitably leads to fatty liver disease. Understanding this pathway opens avenues for novel therapeutic interventions targeting stress-related metabolic disorders.

## INTRODUCTION

Obesity has reached pandemic levels globally (Blüher, 2019). Given the strong correlation between obesity and metabolic disease, understanding the causes of this dramatic increase is crucial for effective prevention and treatment. While excessive consumption of high-fat diets is often considered a primary driver of the obesity epidemic, substantial evidence points to stress and disrupted sleep as significant, yet understudied, contributing factors (Broussard and Van Cauter, 2016; Chaput et al., 2023).

A shared mechanism through which chronic stress and dysregulated sleep act is by disrupting the secretory pattern of glucocorticoids (GCs), essential hormones that control whole body metabolism (Vegiopoulos and Herzig, 2007). In healthy humans, GCs are secreted in circadian oscillations that peak around 7AM and drop low at night (Weitzman et al., 1971). Chronic stressors, such as work-life imbalance and poor sleep quality, “flatten” this rhythm by elevating nighttime troughs and reducing daytime peaks, while maintaining the same average circulating GC levels (Meijer et al., 2023; Bahrami-Nejad et al., 2018) (Figure 1A). Previous studies demonstrated that flattening GC circadian rhythms in mice resulted in large increases in adipose tissue mass, independent of increased food intake or decreased energy expenditure (Bahrami-Nejad et al., 2018; Tholen et al., 2022; Luijten et al., 2019). The link between GC-flattening and obesity is also supported by human studies (Broussard and Van Cauter, 2016; Meijer et al., 2023; Joseph and Golden, 2017).

**Figure 1.**
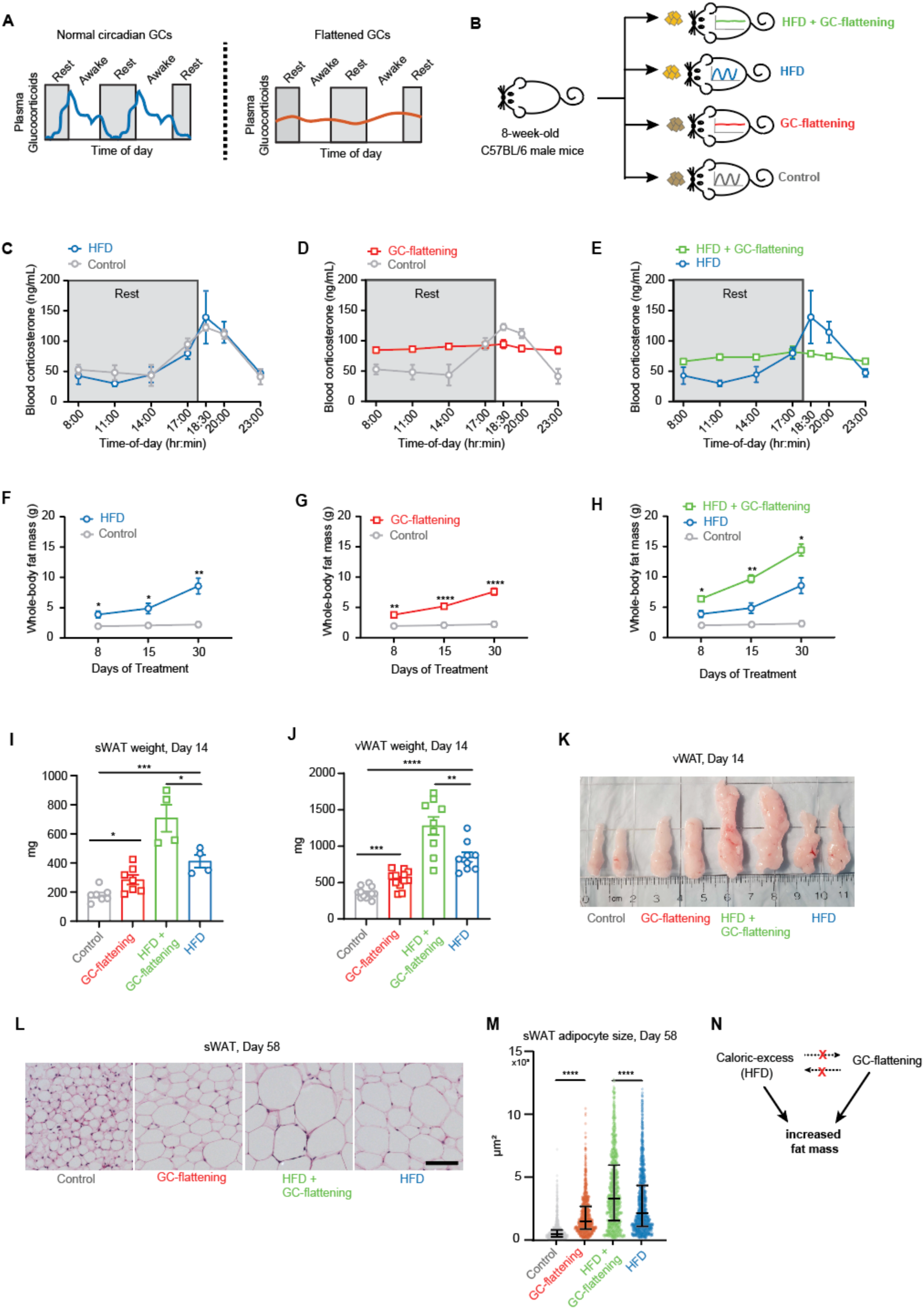
GC-flattening and high-fat diet (HFD) have additive effects in increasing fat mass. (A) Glucocorticoids (GCs) are normally secreted in circadian rhythms in healthy mammals, peaking at the start of the daily awake period and dropping low during the daily rest period. Chronic stressors and dysregulated sleep cause “GC-flattening”, meaning that daily troughs become elevated and daily peaks are reduced, but the average circulating GC levels remain the same. (B) At Day 0, corticosterone pellets to flatten GCs, or placebo pellets, were implanted into 8-week-old C57BL/6 male mice, and the mice were fed either a chow or HFD. The mice were thus divided into four treatment groups: placebo pellet + chow diet (Control), corticosterone pellet + chow diet (GC-flattening), placebo pellet + HFD (HFD); and HFD + corticosterone pellet (HFD + GC-flattening). (C-E) Time course of average corticosterone levels in the mice at Day 58 of treatment. n = 5 mice per group. (F-H) Longitudinal EchoMRI measurements were taken at Days 8, 15, and 30 of treatment. n=7 mice per group, means ± SEM. (I) sWAT fat pad weights at Day 14, n=4-7 mice per group. (J) vWAT fat pad weights at Day 14, n=9-12 mice per group. (K) Representative vWAT fat pads at Day 14. (L) Representative H&E images of sWAT depots at Day 58. Scale bar = 100 µm. (M) Scatter plot showing the size distribution of individual sWAT adipocytes at Day 58 with median and 25th and 75^th^ percentiles marked. n= 6 mice per group. 1000-2000 total number of cells analyzed per group. (N) Scheme explaining how GC-flattening and HFD work through distinct, separable mechanisms to increase fat mass. (F-H) Two-way repeated measures ANOVA with Šídák’s test for multiple comparisons. (I,J) Unpaired t-test. (M) One-way ANOVA with Tukey’s post-hoc test for multiple comparisons. *p < 0.05, **p < 0.01, ***p < 0.001, ****p < 0.0001, ns – not significant.

The dramatic fat mass increase caused by flattening of GC oscillations has so far only been tested in mice fed a low-fat chow diet (Bahrami-Nejad et al., 2018; Tholen et al., 2022; Luijten et al., 2019; Kroon et al., 2021). However, in contemporary society, stress-mediated GC-flattening frequently occurs in the background of a high-fat diet (HFD). Both risk factors are relevant, as current lifestyles generally involve the consumption of processed foods rich in fat (Imamura et al., 2015; Micha et al., 2014; Monteiro et al., 2013; Popkin et al., 2012) and are accompanied by pervasive stress and irregular sleep schedules that disrupt the normal circadian GC rhythms (Koch et al., 2017; Almeida et al., 2023; Spiegel et al., 1999; Miller et al., 2007). While both flattened glucocorticoid (GC) rhythms and a high-fat diet (HFD) are recognized as important contributors to obesity, it is unknown if these two risk factors induce obesity via the same or different pathways. Determining the distinct mechanisms by which they promote obesity would enable the development of more targeted and personalized therapeutic strategies.

Here we induced obesity in mice by flattening the daily GC rhythms or feeding them a HFD. We found that HFD and GC-flattening caused similar increases in fat mass, and these increases were additive. Markedly, both HFD and GC-flattening increased fasting insulin levels but surprisingly, the insulin levels in GC-flattened mice were much higher. We show that this high insulin level increased lipid accumulation in adipose tissue by suppressing lipolysis and by causing a selective insulin resistance in muscle relative to adipocytes that shifted the relative net glucose uptake from muscle to fat. Strikingly, despite the several-fold increase in lipid accumulation in adipose tissue, the liver in GC-flattened mice remained insulin sensitive, free of lipid accumulation (steatosis), and suppressed gluconeogenesis similarly to control mice. We demonstrate that the absence of lipid accumulation in the liver is due to surprisingly normal liver metabolism. Specifically, the livers of GC-flattened mice exhibited typical levels of VLDL secretion and hepatic glucose output, and circulating non-esterified fatty acid (NEFA) levels remained unchanged during fasting. Thus, even though the two risk factors can increase fat mass to similar degrees, we show that only HFD, but not GC-flattening, causes nonalcoholic fatty liver disease due to differences in the liver metabolism and fasting glucose and lipid levels in GC-flattened versus HFD fed mice.

## RESULTS

### GC-flattening and high-fat diet (HFD) have additive effects in increasing fat mass

We first determined how GC-flattening and high-fat diet (HFD) change fat mass over time. To flatten circadian GC rhythms, we implanted mice with pellets that slowly released a low and constant dose of corticosterone, the main physiological GC in mice (Sapolsky et al., 1986), over several weeks. We chose the dose of corticosterone to be released such that the average combined level of circulating GCs (pellet-released plus endogenous corticosterone) was maintained at normal physiological levels (Bahrami-Nejad et al., 2018; Hodes et al., 2012). This pellet protocol recapitulates chronic stress conditions by increasing the daily trough GC level during the rest period and indirectly reducing the peak GC level during the wake period (Bahrami-Nejad et al., 2018; Meijer et al., 2023; Dallman et al., 2000).

To compare the separate and combined effects of GC flattening and HFD, we divided 8-week-old C57/Bl6 male mice into four treatment groups in which mice were (1) fed a chow diet and implanted with a placebo pellet (Control), (2) fed a chow diet and implanted with a corticosterone pellet to flatten GC-rhythms (GC-flattening), (3) fed a high-fat diet and implanted with a placebo pellet (HFD), or (4) fed a high-fat diet and implanted with a corticosterone pellet (HFD + GC-flattening) (Figure 1B). We treated mice for up to 8 weeks. Notably, we verified that high-fat diet (HFD) alone did not alter circadian GC rhythms (Figure 1C). In contrast, GC pellet implantation flattened the circadian GC rhythm in both control and HFD-fed mice (Figure 1D, 1E).

We employed EchoMRI, a non-invasive technique, to track whole-body fat mass in mice over time. We conducted a time-course analysis to assess changes in fat mass across four treatment groups after 8, 15, and 30 days. As anticipated, a high-fat diet (HFD) resulted in a gradual 3-fold increase in fat mass compared to controls (Figure 1F). GC-flattening alone led to a similar, 2.5-fold increase (Figure 1G). Remarkably, the combination of HFD and GC-flattening produced an additive effect, leading to a nearly 5.5-fold increase in fat mass relative to controls (Figure 1H).

Since EchoMRI provides whole-body fat mass measurements without differentiating between specific tissues, we dissected the treated mice to examine changes in individual adipose tissue depots. After 14 days, both subcutaneous white adipose tissue (sWAT) and visceral white adipose tissue (vWAT) depots in GC-flattened, chow-fed mice showed significantly increased mass compared to controls (Figure 1I-J, red versus grey bars). As expected, HFD also significantly increased sWAT and vWAT tissue mass (Figure 1I-J, blue versus grey bars). The combination of GC-flattening and HFD treatment resulted in a remarkable additive increase in both sWAT and vWAT fat mass (Figure 1I-J, green bars; Figure 1K, S1a). By 60 days, sWAT mass in GC-flattened mice was 4-fold higher than controls, with a further additive increase when combined with HFD (Figure S1B-C). In contrast, the additive increase in vWAT plateaued by 60 days (Figure S1D-E).

A common feature of HFD-induced obesity is adipocyte hypertrophy, or increase in the size of adipocytes. To determine whether GC-flattening also induced adipocyte hypertrophy, we next compared adipocyte size across treatment groups by analyzing paraffin-embedded sWAT sections stained with hematoxylin and eosin (H&E). As shown in a representative example of sWAT (Figure 1L), adipocyte size greatly increased after both GC-flattening or HFD, with a further increase when the two treatments were combined. A statistical analysis of the size differences of individual sWAT cells is shown in (Figure 1M). We observed similar additive effects on adipocyte hypertrophy in vWAT (Figure S2A-B).

Both GC-flattening or HFD-feeding increased adipocyte size and total fat mass in mice to similar degrees. However, HFD increased fat mass in mice without altering circadian GC rhythms (Figure 1C and 1F), while GC-flattening increased fat mass in mice on a normal chow diet, independent of caloric excess (Figure 1D and 1G). Taken together, these results support that GC-flattening and HFD independently and additively promote obesity (Figure 1N). Within only 30 days, the combined treatment caused a several-fold increase in adipocyte size and total fat mass, significantly greater than driven by either GC-flattening or HFD alone.

### GC flattening, but not HFD, causes a loss of lean mass

We found that mice fed a high-fat diet (HFD) were heavier than those subjected to GC-flattening for the same treatment time. As expected, HFD led to a continuous increase in body weight (Figure 2A, S3). In contrast, GC-flattening increased the weight at a slower rate, while combining both treatments resulted in an additive increase in weight. Whereas weight increased immediately and continuously in the HFD-mice, the increase in weight in the GC-flattened mice was also delayed relative to control mice and only increased after two weeks (Figure 2A, S3).

**Figure 2.**
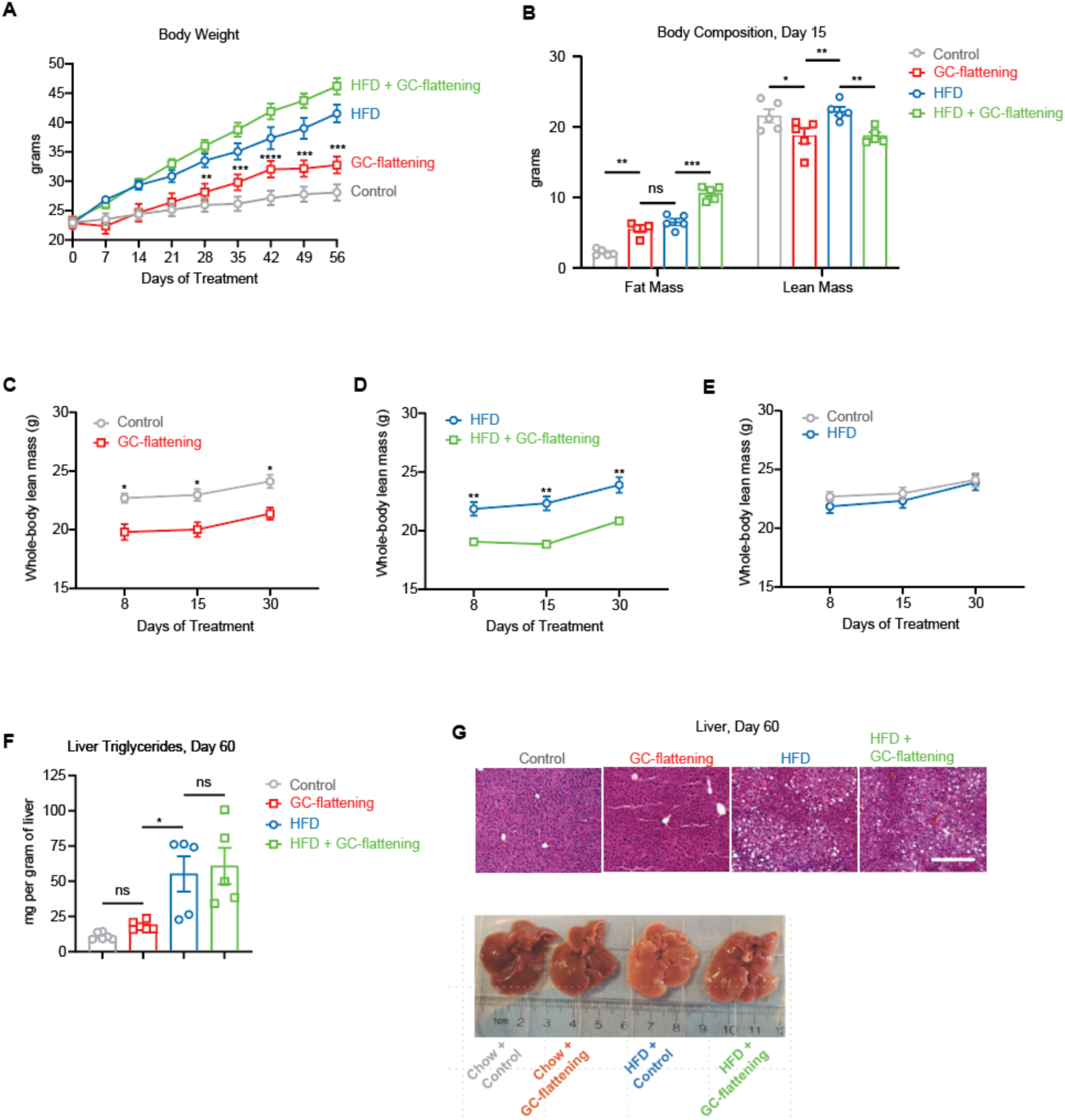
Only GC-flattening resulted in a loss of lean mass, and only HFD caused lipid accumulation in the liver. (A) Time course of body weights of mice in the four treatment groups, n = 5 mice per group. (B) EchoMRI measurements at Day 15, n=5 mice per group. (C-E) Longitudinal EchoMRI measurements of lean mass taken at Day 8, 15, and 30 days of treatment n=7 mice per group. (F) Triglycerides measured in liver lysates from mice at Day 60. n = 5-6 per group, means ± SEMs. (G) Representative liver hematoxylin and eosin (H&E) staining (scale bar = 100 μm) and images. (A-E) Two-way repeated measures ANOVA followed by Šídák’s test for multiple comparisons. In (A) significance values are derived from statistical analysis performed on the differences in body weights for each animal at every time point calculated with respect to its initial body weight as presented in Figure S3. (F) One-way ANOVA with Tukey’s test for multiple comparisons; *p < 0.05, **p < 0.01, ***p < 0.001, ****p < 0.0001, ns – not significant. All data are mean ± SEM.

The delay and slower rate of body weight increase in the GC-flattened versus HFD-fed mice were surprising since the EchoMRI data showed that fat mass in both GC-flattened and HFD-mice increased rapidly and continuously, with a 2.5-fold and 3-fold increase in fat mass respectively after 30 days (Figure 1F-G). We thus considered that a parallel lean (muscle) mass change could explain the slower, delayed body weight gain in GC-flattened versus HFD-fed mice. Using again EchoMRI body composition measurements, we indeed found a marked reduction in lean mass in the GC-flattened mice after 15 days. In contrast, the HFD-fed mice did not lose lean mass, and their weight increase was mostly driven by increased fat mass (Figure 2B). Moreover, this reduction in lean mass was also observed in mice in subjected to a combined HFD-fed and GC-flattened treatment (green bar, Figure 2B). The selective effect of GC-flattening, but not of HFD, on reducing lean mass can also be seen in a time course analysis (Figure 2C-E). We conclude that a primary consequence of HFD is an increase in fat mass at a constant lean mass, resulting in a large increase in overall body weight. In contrast, a primary consequence of GC-flattening is an increase in fat mass concurrent with a loss in lean mass, resulting in only a small increase of overall body weight.

### Unlike HFD, GC-flattening does not cause fat accumulation in the liver

A major clinical consequence of a high-fat diet (HFD) is the accumulation of fat in the liver that can lead to non-alcoholic liver disease (NAFLD) (Geißler et al., 2022). Previous studies showed that GC treatments can also lead to fat accumulation in the liver (Campbell et al., 2011). However, the doses of GCs administered in these previous studies were much higher than endogenous levels of GCs, and thus well above the continuously administered GC doses used in the current study which kept the average circulating GC levels at normal levels, more faithfully replicating physiologically-relevant chronic stress conditions (Bahrami-Nejad et al., 2018; Meijer et al., 2023; Broussard and Van Cauter, 2016). Strikingly, with GC-flattening, no increase in the fat content of the liver was observed, even after 60-days (Figure 2F). Moreover, whereas GC-flattening on top of a HFD massively increased adipose tissue mass (i.e. Figure 1K), it did not increase hepatic lipid accumulation beyond that induced by the HFD alone (Figure 2F).

These findings are supported by H&E staining, which showed no increase in lipid deposition in the livers of GC-flattened compared to control mice. In contrast, HFD greatly increased lipid accumulation in the liver (Figure 2G, top). Moreover, representative gross images showed that HFD-fed mouse livers were paler than GC-flattened mouse livers (which looked similar to control mice), again indicating that HFD, but not GC-flattening leads to lipid accumulation in the liver (Figure 2G, bottom). Taken together, these results selectively link HFD -but not GC-flattening - to non-alcoholic fatty liver disease (NAFLD). The lack of liver fat accumulation in GC-flattened mice was unexpected since the size of the sWAT and vWAT fat depots both increased (Figure 1I-K), raising the questions how GC flattening causes an increase in fat mass while losing lean mass and why there is no fat accumulation in the liver.

### GC-flattening rapidly drives hyperinsulinemia to much higher levels than HFD

Increased fat mass in humans and rodents is associated with high fasting insulin and glucose levels due to increased insulin resistance (Odeleye et al., 1997; Corkey, 2012; Janssen, 2021). We thus compared the fasting insulin and glucose levels for the 4 different treatment groups. By 14 days, the HFD-fed mice showed the expected increases in fasting insulin (Turner et al., 2013), increasing to an approximately 2-fold increase in fasting insulin after seven weeks of HFD feeding (Figure 3A). In contrast, after only three days of treatment, GC-flattening increased fasting insulin to higher levels, and these levels remained more than 6-fold higher than the control mice and 3-fold higher than the HFD-fed mice for the duration of treatment (Figure 3B). Combining HFD and GC-flattening caused even higher persistent increases in fasting insulin levels (Figure 3C).

**Figure 3.**
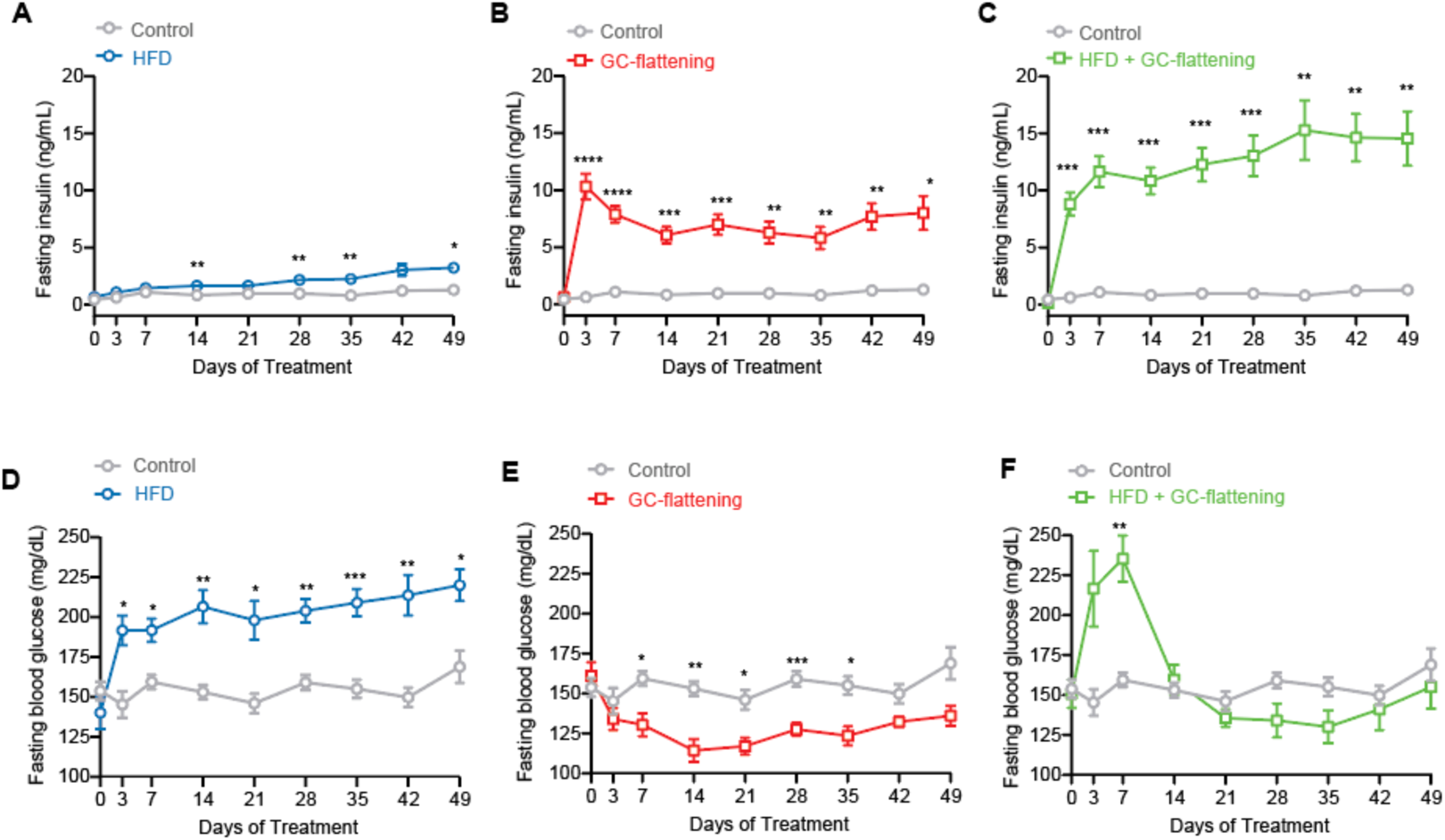
GC-flattening rapidly generated fasting insulin levels several-fold higher than the insulin increase from HFD, and without elevated circulating glucose levels. (A-C) Fasting insulin and (D-F) glucose levels were measured over 7 weeks in control or GC-flattened mice fed a chow or HFD. n=10 mice per group across two experimental repeats. Two-way repeated measures ANOVA followed by Šídák’s test for multiple comparisons. *p < 0.05, **p < 0.01, ***p < 0.001, ****p < 0.0001, ns – not significant. All data are means ± SEM.

The same time course analysis for glucose showed the expected HFD-mediated increase in the fasting glucose levels over control mice (Turner et al., 2013)(Figure 3D). However, GC-flattening had the opposite effect, reducing fasting glucose to a level below that of control mice (Figure 3E). Combining GC-flattening and HFD resulted in an initial increase in fasting glucose levels but again ended with glucose levels similarly low as in the control mice, much lower than the level in HFD-fed mice (Figure 3F). That GC-flattened mice have fasting glucose levels lower than control animals was surprising since hyperinsulinemia is generally believed to result from increased fasting glucose levels. Taken together, both GC-flattening and HFD increase fasting insulin levels but the crucial difference between the two is that HFD-fed mice have much higher, while GC-flattened mice have lower, fasting glucose levels than control mice.

### Glucose tolerance in GC-flattened mice due to high insulin and a higher insulin setpoint

To understand the metabolic process controlling the liver and adipose tissue in GC-flattened mice, we carried out glucose and insulin tolerance tests (GTT and ITT) in mice after GC-flattening, HFD, or both treatments combined. We chose to carry out the measurements after 8 weeks to allow enough time for the effects of HFD feeding to manifest (Geißler et al., 2022). As shown by the grey line in Figure 4A, injected glucose is cleared in approximately 120 minutes in chow-fed, control mice. Markedly, injected glucose cleared faster in GC-flattened mice compared to the control mice (Figure 4A) likely due to increased insulin secreted in response to injected glucose as we have previously observed (Tholen et al., 2022), while HFD-fed mice had the expected worse glucose clearance than control mice. Moreover, GC-flattening on top of a HFD resulted in improved glucose clearance (Figure 4A). Thus, GC-flattening improved glucose tolerance in both control and HFD mice.

**Figure 4.**
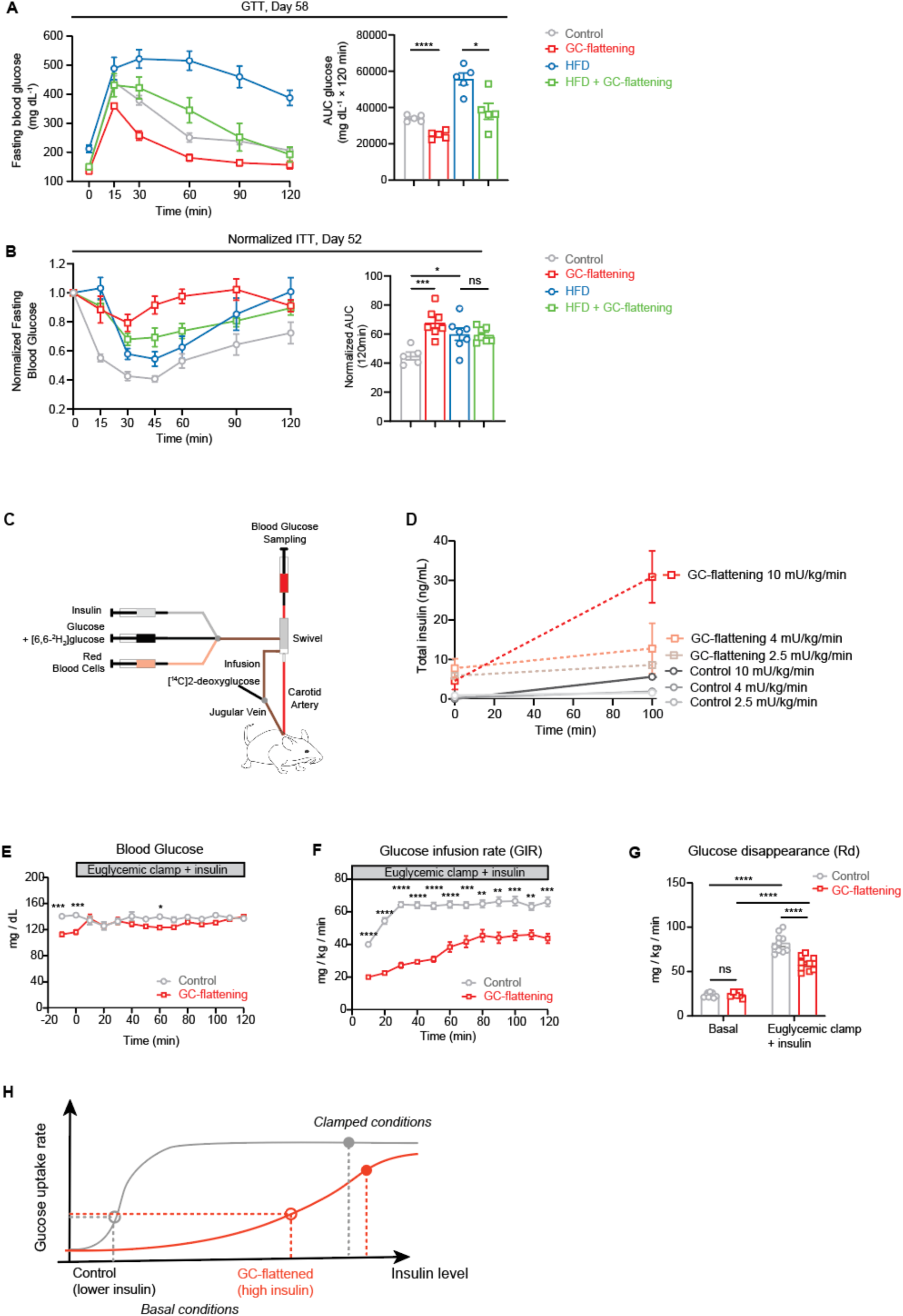
Glucose tolerance in GC-flattened mice due to a higher insulin setpoint. (A) An intraperitoneal glucose tolerance test (GTT) was performed on control or GC-flattened mice fed a chow or HFD following 58 days of treatment. The panel on the right shows the cumulative area under the curve during the GTT. n=5 mice per group. (B) An intraperitoneal insulin tolerance test (ITT) was performed following 52 days of treatment. Blood glucose levels measured during the ITT were normalized to starting blood glucose levels in each treatment group. The panel on the right shows the cumulative area under the normalized curve during the ITT. n=5-7 mice per group. (C-G) Hyperinsulinemic-euglycemic clamp measurements were performed on chow-fed control or GC-flattened mice at 14 days of treatment. n=8-11 mice per condition. (C) Mice were implanted with catheters in both the jugular vein and a carotid artery for infusions and sampling, respectively. Arterial glucose was clamped at 140 mg/dl for a total of 155 min with a continuous infusion of [6,6-2H2]glucose and fixed amount of insulin. Mice also received heparinized saline-washed erythrocytes from donors during the clamp to prevent reduction in hematocrit. For tissue specific glucose uptake measurements mice received an intravenous bolus of radioactive 2[14C]deoxyglucose at 120 min and sacrificed for tissue dissections 35 min later. (D) Control and GC-flattened mice were injected with increasing doses of insulin as indicated in a pilot experiment to determine the rate at which insulin must be infused for the hyperinsulinemic-glycemic clamp measurements. (E) Blood glucose levels (infused glucose + endogenous glucose) was monitored throughout the clamp at 10-minute intervals by sampling from the arterial catheter. Blood glucose was maintained at euglycemia (∼140 mg/dL). (F) Glucose infusion rate (GIR) in the venous catheter needed to maintain euglycemia. (G) Rate of glucose disappearance from the circulation (Rd), an indicator of whole-body glucose uptake was measured from blood collected from mice 10 min prior (Basal) to clamping and as an average of blood collected at four different timepoints between 80-120 minutes after the clamp was initiated (Euglycemic clamp+insulin). (H) Schematic depicting the increased insulin setpoint in GC-flattened mice. (A,B) Unpaired t-test. (C,E) Mixed-effects model (REML) followed by Šídák’s post-hoc test for multiple comparisons. *p < 0.05, **p < 0.01, ***p < 0.001, ****p < 0.0001, ns – not significant. All data are mean ± SEM.

We next carried out an insulin tolerance test (ITT), where insulin is injected into the mice rather than glucose. As shown by the grey line in Figure 4B, glucose levels in control mice rapidly fell in the first 30 minutes following injection of insulin, and the HFD-fed mice showed the expected slower drop in circulating glucose levels. However, despite being more glucose tolerant, chow-fed and HFD-fed GC-flattened mice did not lower glucose as effectively as control mice after insulin injection (Fig 4B, S4). This is not surprising since fasting insulin levels are already high in GC-flattened mice before insulin injections, and additional insulin is not expected to further activate the receptor.

Our interpretation was that GC-flattened mice are not insulin resistant since the already high insulin level is effectively disposing of increased glucose with added insulin having only minimal effect. Also, the ITT is inherently limited due to its single-timepoint insulin administration, also neglecting time-dependent mechanisms such as hepatic glucose production, which can modulate glucose levels independently of tissue glucose uptake. To investigate whether the insulin sensitivity is altered in GC-flattened mice, we therefore utilized hyperinsulinemic-euglycemic clamp (”insulin-clamp”) studies, which enable constant insulin levels to be maintained in the mice throughout the experimental period (Alquier and Poitout, 2018) (Figures 4C, S5).

Because the GC-flattened mice have much higher circulating insulin levels compared to control mice (Figure 3B), we first determined the rate at which insulin had to be infused into the control and GC-flattened mice to obtain similar serum insulin levels. We infused 3 different doses of insulin (2.5, 4, and 10 mU/kg/min) and compared the resulting total circulating insulin levels (sum of the infused plus endogenous circulating insulin; Figure 4D). We used an infusion rate of 2.5 mU/kg/min for the GC-flattened mice and 10 mU/kg/min for the control mice since these doses generated similar steady-state levels of total insulin in the blood.

After clamping the insulin infusion rates, we next infused glucose at rates needed to maintain normal total glucose levels (∼140 mg/dL) (Figure 4E). Lower glucose infusion rate (GIR) corresponds to increased insulin resistance, and the GIR in the GC-flattened mice was approximately half that of the control mice (Figure 4F). The Rd value reflects how fast glucose disappears from the bloodstream and thus corresponds directly to how fast glucose is taken up into tissues. However, as shown in Figure 4G, despite showing insulin-resistance, the GC-flattened mice are still able to uptake glucose effectively, as shown by the increased glucose disappearance (Rd) when insulin is increased between basal and clamped conditions.

The combined results from the glucose tolerance test (GTT), insulin tolerance test (ITT), and insulin-clamp experiments point to a ‘shift in the insulin setpoint’. Figure 4H depicts this concept, showing that GC-flattened mice have much higher basal insulin levels (around 8-fold greater, as measured in Figure 3B) but show increased glucose clearance as measured by the GTT assay where both glucose and secreted insulin levels change. However, GC-flattened mice show similar glucose uptake rates as control mice when glucose is infused to maintain euglycemia during the insulin clamp, as depicted in the model (Figure 4H). Furthermore, despite their pronounced hyperinsulinemia, GC-flattened mice can robustly glucose uptake in response to increased insulin (“clamped conditions”, Figure 4H). Notably, to achieve similar blood insulin levels in control and GC-flattened mice during the clamp experiment, a four-fold higher insulin infusion rate was used for the control mice (10 mU/kg/min versus 2.5 mU/kg/min in the GC-flattened mice, as shown in Figure 4D). We expect that glucose uptake rates in GC-flattened mice would have reached similar levels as in control mice if more insulin had been also infused into the GC-flattened mice. Therefore, as depicted in the model presented in Figure 4H, these observations suggest that GC flattening induces a shift in the insulin setpoint. GC-flattened mice remain responsive to insulin and are capable of controlling glucose uptake, but only at a considerably elevated relative insulin level or ‘insulin setpoint’.

### In contrast to HFD, GC-flattening preserves insulin sensitivity of the liver

Surprisingly, we found that the liver remained insulin-sensitive in GC-flattened mice, as evidenced by there being no significant difference from control mice in hepatic glucose output in both basal and insulin-clamped conditions (Figure 5A). Western blot analysis confirmed a downregulation of the liver phosphoAKT levels in HFD-fed mice compared to control mice, but without a significant difference in GC-flattened mice (Figure 5B).

**Figure 5.**
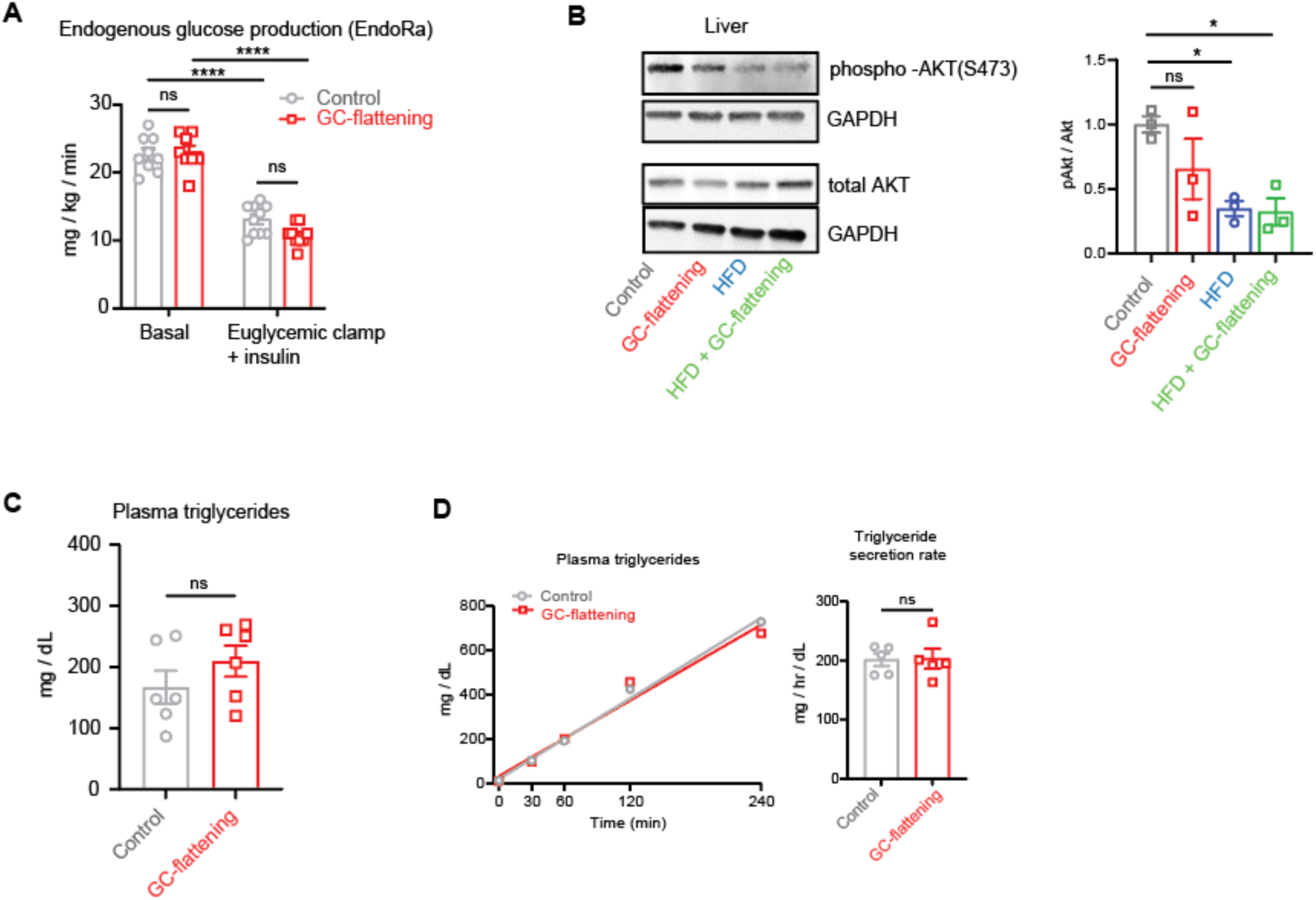
GC-flattened mice preserve liver sensitivity and metabolic function. (A) The rate of endogenous glucose production (EndoRa), an indirect measure of hepatic glucose production was measured from blood collected from mice 10 min prior (Basal) to clamping and as an average of blood collected at four different timepoints between 80-120 minutes after the clamp was initiated (Euglycemic clamp+insulin). (B) Western blot images (left) and quantification (right) of phospho-Akt Ser473, Total Akt and GAPDH (loading control) levels in mouse livers 60 days after pellets were implanted. Mice were fasted for 4-6h and received an insulin injection (0.75 UI Insulin/kg BW) 10 min before they were sacrificed for tissue collection. n=3 mice per condition. (C) Triglycerides were measured from blood plasma of control and GC-flattened mice on Day 60 of treatment. n=6 mice per condition. (D) Tyloxapol was injected into fasted mice at Day 14 of treatment to inhibit the uptake of circulating VLDL into tissues. n=5 mice per condition. *Left,* Triglyceride levels in blood were measured at the indicated times *Right,* Triglyceride secretion rates were calculated as the slope of the line for each condition. (B, C, D) Unpaired t-test was used to compare for statistical differences between indicated treatment groups, *p<0.05, ns -not significant.

Previous work has also shown that a critical consequence of insulin resistance of the liver is increased secretion of triglycerides VLDL particles (Schweiger et al., 2017). Consistent with GC-flattened mice retaining liver insulin sensitivity, we found that triglyceride levels in the blood of GC-flattened mice were indistinguishable from control mice (Figure 5C). It was nevertheless conceivable that even though the levels were the same, the triglyceride flux (both the synthesis and disposal of VLDL) might be increased in GC-flattened mice. We thus directly determined whether the rate of release of triglycerides is different in GC-flattened and control mice by injecting tyloxapol, a small molecule that blocks the uptake of triglycerides. We measured the accumulating triglycerides in the blood of fasted mice just before the injection (t = 0), and at 30, 60, 120, and 240 minutes after injection. As shown in Figure 5D, the rate of triglyceride/VLDL secretion from the liver was the same in GC-flattened and control mice, demonstrating that the liver’s handling not only of glucose but also of lipids is overall normal in GC-flattened mice.

Taken together, our results demonstrate that GC-flattening induces insulin resistance in muscle without that the liver loses its insulin sensitivity. This represents a unique GC-flattened metabolic state that is distinct from the HFD state where all three peripheral tissues are insulin resistant and there is fatty liver disease (Geißler et al., 2022; Lang et al., 2019; Surwit et al., 1988).

### Adipose mass increases due to suppression of lipolysis and a relative shift of glucose uptake from muscle to fat

The lack of lipid accumulation in the liver, despite significant increases in adipose tissue mass, made us speculate that the very high hyperinsulinemia in GC-flattened mice could be suppressing lipolysis in the adipose tissue and preventing release of non-esterified fatty acids (NEFAs), which are a major source of lipid normally taken up by the liver (Hodson and Gunn, 2019). In HFD conditions, the insulin-mediated suppression of lipolysis is absent due to the loss of insulin sensitivity of adipose tissues, which results in increased release of non-esterified fatty acids levels and subsequent accumulation in the liver (Lian et al., 2020; Hodson and Gunn, 2019).

To assess whether adipose tissue in GC-flattened mice could still regulate insulin-mediated lipolysis despite displaying insulin resistance to glucose uptake compared to control mice, we measured NEFA levels in mice that had been fasted for 5 hours, before and immediately after raising insulin levels. We found that NEFA levels in the blood of both control and GC-flattened mice were similarly suppressed by insulin, as shown by the similar large drop in NEFA levels between basal and clamp conditions (Figure 6A) and similar percent suppression of plasma NEFAs in each animal (Figure 6B). While basal NEFA levels were slightly higher in GC-flattened mice, this was much less than expected given their 2.5-fold increase in fat mass at the Day 14 measurement timepoint (Figure 1G, 2B), supporting a suppression of basal lipolysis in GC-flattened mice. The observation that GC-flattened mice maintain insulin-mediated suppression of lipolysis indicates their continued responsiveness to insulin. This provides further support for the ‘insulin setpoint’ model (Figure 4H), which posits that the significantly increased insulin levels in GC-flattened mice allow them to maintain glucose disposal capacity despite mild insulin resistance. Moreover, the high insulin levels not only contribute to slightly lower glucose levels in GC-flattened mice compared to controls by suppressing hepatic glucose production and enhancing glucose uptake, but also effectively suppress lipolysis, keeping circulating NEFA levels low, comparable to control mice.

**Figure 6.**
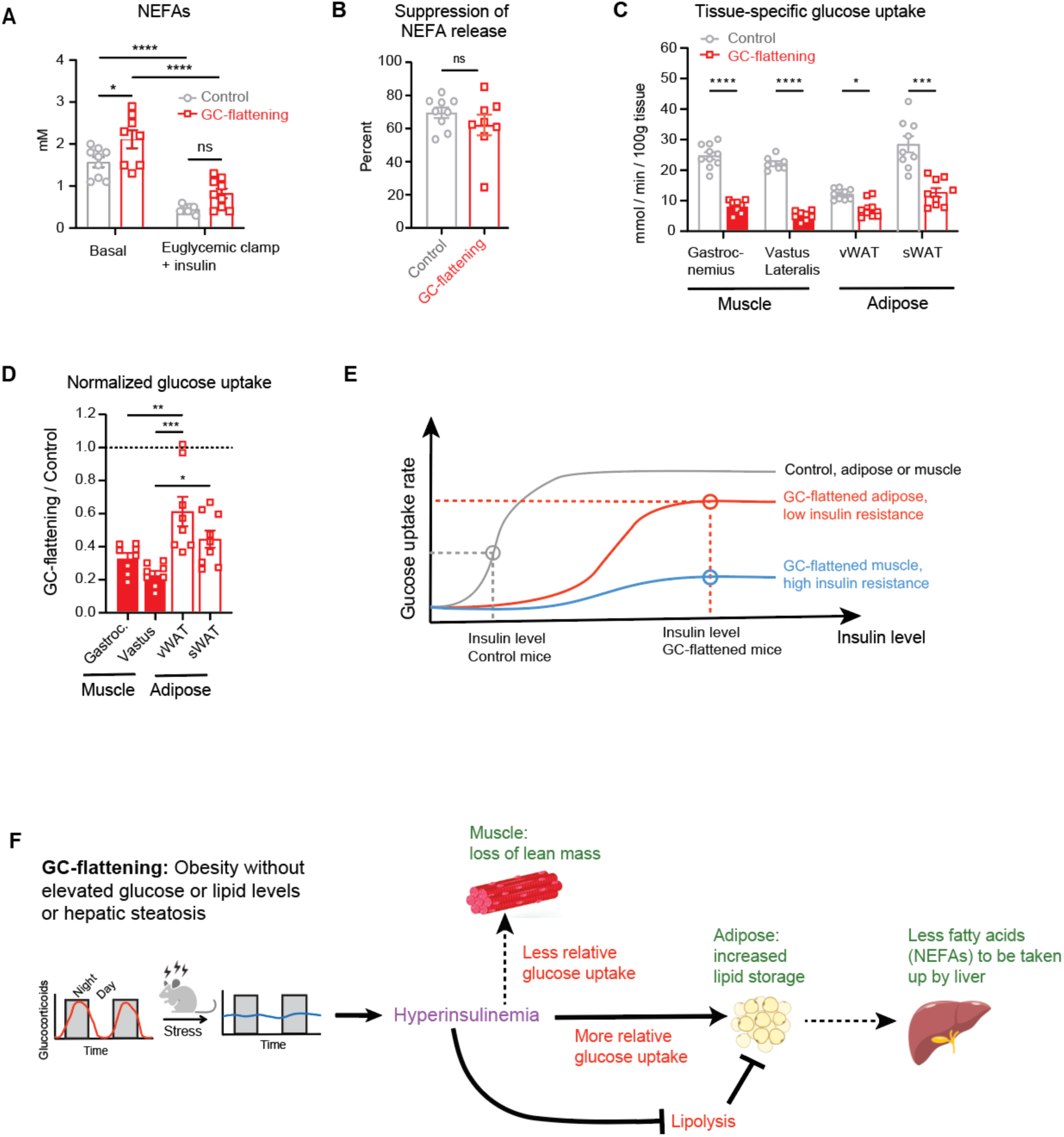
Adipose mass increases due to suppression of lipolysis and a relative shift of glucose uptake from muscle to fat. (A) Non-esterified fatty acid (NEFA) levels were measured from the blood of fasted control and GC-flattened mice 10 min prior (Basal) and 145 min after the clamp was initiated (Euglycemic clamp+insulin). Mixed-effects model (REML) followed by Šídák’s post-hoc test for multiple comparisons. *p < 0.05, ****p < 0.0001, ns – not significant. All data are mean ± SEM. (B) The percent suppression in NEFA release was calculated for each mouse as the ratio of its plasma NEFA levels measured 145 minutes after the clamp was initiated divided by its plasma NEFA levels measured 10 min prior to the clamp. Unpaired t-test was used to compare means, ns – not significant. (C) All mice received an intravenous bolus of non-metabolizable glucose [14C]2DG, 120 minutes after clamping was initiated and sacrificed 35 minutes later (total clamping for 155 minutes) to collect the tissues. [14C]2DG uptake was measured from indicated tissues. Mixed-effects model (REML) followed by Šídák’s post-hoc test for multiple comparisons. *p < 0.05, ***p < 0.001, ****p < 0.0001, ns – not significant. All data are mean ± SEM. (D) Glucose uptake into different tissues of GC-flattened mice was ratioed to the glucose uptake in corresponding tissues of control mice. One-way repeated ANOVA with Tukey’s test for multiple comparisons. *p < 0.05, **p < 0.01, ***p < 0.001. All data are mean ± SEM. (E) In GC-flattened mice, muscle tissue is more insulin resistant than adipose tissue. The very high hyperinsulinemia in GC-flattened mice allows adipose tissue to still uptake glucose at basal conditions, even more than in control mice in basal conditions. (F) Schematics of the mechanisms driving significant increases in adipose tissue mass in response to GC-flattening. GC-flattening and the resulting hyperinsulinemia cause a shift in the relative amounts of glucose taken up by muscle and adipose tissue, as well as cause increased lipid storage in adipose tissue due to suppression of lipolysis and increased de novo lipogenesis and lipid uptake. These effects combine to result in a reduction in lean mass and a selective increase in the fat mass (obesity) without hepatic steatosis or a significant increase in circulating glucose and lipid levels such as occurs in HFD-driven obesity.

We next sought to determine whether tissue-specific differences in insulin-mediated glucose uptake could account for the paradoxical increase in adipose tissue and reduction in lean tissue observed in GC-flattened mice. To this end, we measured incorporation of radiolabeled glucose into muscle and fat tissues while insulin levels were clamped at similar levels in control and GC-flattened mice. We found that insulin-stimulated glucose uptake was reduced in both muscle and adipose tissue of GC-flattened mice (Figure 6C, S6), consistent with both types of tissue being insulin resistant. However, the GC-flattened mice could still robustly uptake glucose in response to insulin, as demonstrated by the GTT results (Figure 4A) and from calculating the rate of disappearance of glucose from the bloodstream in the insulin-clamp experiments (Figure 4G).

In these insulin-clamped conditions, a marked difference was observed: GC-flattened mice demonstrated a disproportionately higher glucose uptake into fat tissue compared to muscle, when contrasted with control mice (Figure 6D). This observation indicates that a greater level of insulin resistance in muscle relative to adipose tissue could provide a mechanistic explanation for the increased fat mass and decreased lean mass resulting from GC-flattening. Thus, the concept developed in Figure 4H that the insulin sensitivity setpoint is shifted in GC-flattened mice, allowing them to maintain high glucose uptake rates despite obesity, can be expanded upon by adding that these mice have a higher loss of insulin sensitivity in muscle relative to fat (Figure 6E).

## DISCUSSION

Our study investigated the mechanisms by which GC-flattening and a high-fat diet (HFD) induce obesity. Both conditions increased insulin levels and resulted in a substantial and additive increase in fat mass, but by different mechanisms. Insulin tolerance tests (ITTs) suggest insulin resistance in GC-flattened mice. However, glucose tolerance tests (GTTs) and hyperinsulinemic-euglycemic clamp experiments reveal improved glucose clearance compared to controls, even more so than in HFD-fed mice. This paradox is resolved by understanding that GC-flattened mice exhibit significantly higher insulin levels, compensating for reduced insulin sensitivity in adipose tissue and liver. Rather than having the canonical insulin resistance in which adipose, liver, and muscle tissues are all dysfunctional and fail to respond to insulin, we demonstrated that adipose tissue and liver of GC-flattened mice can effectively respond to insulin but require higher insulin levels. Overall, our findings argue that adipose tissue and liver of GC-flattened mice have a shift in their “insulin setpoint” without losing their ability to respond to insulin. In contrast, muscle has a higher degree of insulin resistance relative to adipose tissue, explaining the relative increase in glucose uptake in adipose tissue than muscle and the reduction in lean mass in GC-flattened mice (Figure 6F).

Our discovery of a higher “insulin setpoint” in GC-flattened mice is supported by there being no impairment in liver metabolic function, despite very high levels of hyperinsulinemia. Hepatic glucose output was equally suppressed in GC-flattened and control mice, indicating normal liver insulin sensitivity (Figure 4E). The livers of GC-flattened mice had normal triglyceride secretion and no hepatic steatosis. However, insulin-clamp experiments confirm reduced insulin-stimulated glucose uptake in both muscle and adipose tissues of GC-flattened mice compared to controls (Figure 4F, 6C). Thus, GC-flattening induces insulin resistance in muscle and adipose tissues while preserving liver insulin sensitivity, a pattern distinct from HFD-induced obesity, where all three tissues develop insulin resistance (Geißler et al., 2022; Lang et al., 2019; Surwit et al., 1988). We conclude that GC-flattening-induced obesity preserves liver function, a key distinction from HFD-induced obesity which leads to hepatic steatosis.

We observed that fasting basal non-esterified fatty acid (NEFA) levels were not elevated in GC-flattened mice, despite their significantly increased fat mass. This indicates that elevated insulin levels suppress lipolysis in adipose tissue, thereby inhibiting fatty acid release during fasting and promoting triglyceride accumulation. Moreover, our prior RNA-seq analysis of adipose tissue from GC-flattened mice revealed a substantial upregulation of genes associated with glucose uptake, lipid uptake, and lipogenesis (Tholen et al., 2022). Thus, the adipose tissue in GC-flattened mice remains insulin responsive and can grow in size through increased glucose and fatty acid uptake, as well as de novo lipogenesis and suppression of lipolysis.

In contrast to HFD-fed mice, GC-flattened mice displayed a concomitant loss of lean mass with the increase in fat mass. We found that GC-flattening led to more pronounced insulin resistance in muscle compared to fat, explaining how a combination of a shift of glucose uptake from muscle to fat and a still functioning fatty acid uptake can generate a parallel fat mass increase and lean mass loss. This lean mass reduction was also evident when GC-flattening and HFD were combined. Thus, GC-flattening-induced obesity is driven by increased glucose uptake into fat relative to muscle, coupled with enhanced insulin-mediated lipid synthesis and storage in adipose tissue, resulting in a unique shift from lean to fat mass, distinct from high-fat diet-induced obesity (Figure 6F).

Previous studies suggested that increased caloric intake or lowering of thermogenesis are potential alternative explanations for how GCs promote obesity (Peckett et al., 2011). However, in contrast to our protocol, these studies typically used high pharmacological doses of GCs or used synthetically-stabilized GCs (Gasparini et al., 2016; Karatsoreos et al., 2010; Campbell et al., 2011; Harno et al., 2021). Here we used pellets that released corticosterone, the main physiological GC in mice (Sapolsky, 2021), rather than synthetic GCs. Since our goal was to mimic chronic stress by removing the troughs and peaks while keeping the average GC level near physiological levels, we took care to avoid high pharmacological doses and instead used GC doses that maintained physiological mean circulating levels of GCs. Using this GC-flattening protocol, we and others have observed that the increased fat mass is not the result of increased food intake, decreased energy expenditure, or defective thermogenesis (Tholen et al., 2022; Luijten et al., 2019).

In conclusion, our study reveals that two risk factors of obesity, GC-flattening and HFD, can cause massive and additive increases in fat mass. However, they induce distinct metabolic states, characterized by the presence or absence of non-alcoholic fatty liver disease (NAFLD) (Figure 5). The unique metabolic profile observed with GC-flattening can be attributed to maintained glucose control at elevated insulin levels, a phenomenon best described as a shifted insulin sensitivity setpoint rather than insulin resistance. Moreover, our findings provide a mechanistic understanding of how obesity can develop without hepatic lipid accumulation, opening new avenues for the development of NAFLD therapies that are independent of obesity treatment.

## Methods

### Mice and diets

Seven-week-old C57BL6/J male mice were obtained from The Jackson Laboratory (cat. 000664) and housed in the animal facility to acclimate for 7 days prior to the start of experiments. Mice were housed in small groups of five or fewer on a 12h light/dark cycle (lights on at 6:00 h) in the animal facility at Weill Cornell Medicine with free access to drinking water and fed standard chow diet (PicoLab Rodent Diet 20, 5053; Lab Diets, St. Louis, MO) except when food was restricted during fasting. For diet-induced obesity, mice were fed HFD (60% kcal from fat; D12492; Research Diets, New Brunswick, NJ) ad libitum. Ear tags were used to identify individual mice, and researchers were blinded to the treatment groups. Using a method similar to block randomization, experimental and control groups were matched for body weight before being assigned to a treatment group so that the average baseline body weight across groups was equal at the start of the experiment. Experimental and control mice were mixed in the same cages in order to remove cage-to-cage variations. All mice were dissected between 13:00 and 18:00, alternating corticosterone-treated and control mice.

Prior to collection of blood and tissue, mice were fasted for 6 h during the light cycle with free access to water unless stated otherwise. Blood was collected by tail snip or cardiac puncture. Blood was collected in ethylenediaminetetraacetic acid (EDTA) coated tubes, centrifuged (6,000 x *g* for 20 min at 4°C), and the plasma supernatant was then collected and stored at −20°C. Harvested tissue samples were either fixed in buffered zinc formalin fixative (Cancer Diagnostics Inc., 171) and paraffin embedded, or snap-frozen in liquid nitrogen and stored at −80°C. Protocols for animal use and euthanasia were approved by the Institutional Animal Care and Use Committee of Weill Cornell Medical College.

### Corticosterone administration in mice

To flatten circadian glucocorticoid (GC) oscillations, mice were implanted subcutaneously with pellets that released corticosterone for either 21 days or 60 days (21-day release, 5 mg, Catalog number G-111; 60-day release, 15 mg total, Catalog number SG-111; Innovative Research of America, Sarasota, FL, USA). Placebo pellets (Catalog number C-111 or SC-111) were implanted as control. Prior to pellet implantation, mice were anesthetized via continuous inhalation of isoflurane. Once anesthetized, an incision equal in diameter to that of the pellet was made on the lateral side of the neck, and the pellet was inserted 2 cm from the incision site using forceps or a trocar. Mice weighed an average of 24.1 ± 1.2 g, which resulted in a daily dose of 10 mg/kg/day.

### Measurement of corticosterone in blood serum

Blood was taken at multiple time points (8:00, 11:00, 14:00, 17:00, 18:30, 20:00, and 23:00) over a 15 h time period. At the first time point, blood was taken by nicking the tail vain. Blood samples collected at subsequent time points were taken by removing the crust formed after first blood withdrawal. The corticosterone concentration in blood plasma was determined using the Enzyme Immunoassay kit (K014-H1, Arbor Assays, Michigan, USA) following the manufacturer’s instructions.

### Fasting glucose and insulin measurements

Mice were fasted for 6 hours in clean cages with no food, food dust, or feces in the bedding, and sample collection was done between 3-5 PM. Blood was drawn via tail snip, and blood glucose levels was measured with a glucometer (Diathrive, Salt Lake City, Utah). An additional 30 µL of blood for insulin measurements was collected into EDTA-coated tubes for plasma separation. Blood samples collected in subsequent weeks were done by removing the scab formed after the first blood draw. The insulin concentration in blood plasma was determined using the Ultra-Sensitive Mouse Insulin ELISA Kit (Cat. 90080, CrystalChem, Illinois, USA), following the manufacturer’s instructions.

### Tolerance tests

Before the procedure, mice were fasted for 6 h starting at 9AM in clean cages with no food, food dust, or feces in the bedding. For the glucose tolerance test (GTT), mice received 2 g of glucose (Sigma G8270)/kg of body weight by intraperitoneal injection. For the insulin tolerance test (ITT), mice were injected with 0.75 units/kg body weight of human insulin (Humulin R (U-100), Eli Lilly) by intraperitoneal injection. Blood samples to measure glucose levels were collected before injection and 15, 30, 45, 60, and 120 min after injection and analyzed as described above.

### Hepatic triglyceride secretion rates

Rates of hepatic triglyceride secretion were determined in mice that were fasted for 5 hours (Steensels et al., 2020). The lipoprotein lipase inhibitor Tyloxapol (500 mg/kg body weight) (Sigma-Aldrich, St. Louis, MO) was administered by retro-orbital injection during the light cycle. Tail snip blood samples (25 μl) were collected into Eppendorf tubes containing EDTA prior to and after Tyloxapol injection at regular intervals for up to 4 h. Plasma triglyceride (TG) concentrations were determined using enzymatic assays as described below. Rates of hepatic TG secretion were calculated from the time-dependent linear increases in plasma TG concentration following Tyloxapol administration.

### Measurement of circulating concentrations of FFA and triglycerides

Following euthanasia, plasma was collected and stored at −80°C. FFA contents in blood serum were determined using the Non-Esterified Fatty Acid (NEFA) -HR (2) Kit (Fujifilm Wako, 999-34691, 995-34791, 991-34891, 993-35191, 276-76491, and 997-76491) following the manufacturers’ protocols. Triglyceride concentrations in blood serum were measured using the L-Type Triglyceride M Kit (Fujifilm Wako, 994-02891, 990-02991, 992-02892, 998-02992, and 464-01601) following the manufacturer’s instructions.

### Measurement of liver triglycerides

Livers were harvested from the mice dissected between 1-5 pm. The left lateral lobe was immediately snap frozen in liquid nitrogen and stored at -80°C until analysis. 30 mg of frozen liver was digested in 180 µL of fresh alcoholic KOH (2:1 100% ethanol to 30% KOH) in a shaking incubator (Labnet VorTemp 56) at 700 rpm, 60°C for 1 h or until the tissue was completely dissolved. 1 M MgCl_2_ was added to the digest at 1:1.08 ratio to 1 M MgCl_2_ and vortexed. After 10 mins of incubation on ice, the samples were centrifuged at 1400 rpm for 30 mins. The supernatant was collected, and triglyceride content was measured using the Stanbio LiquiColor Triglycerides Kit (Stanbio, 2100), following previously published protocols(Taylor et al., 2021).

### Liver and adipose tissue histology

For subcutaneous white adipose tissue (sWAT), we took the inguinal white adipose tissue. For visceral white adipose tissue (vWAT), we took the epididymal white adipose tissue. Liver, sWAT, and vWAT depots were harvested from mice, sliced into small chunks measuring approximately 5mm x 5mm (liver) and 2mm x 2mm (adipose tissue), placed into plastic cassettes to allow for good penetration of the fixing solution, and fixed for 24h at 4°C in buffered zinc formalin fixative (Cancer Diagnostics Inc., 171). The tissue chunks were then rinsed with PBS and stored in PBS at 4°C until they were embedded in paraffin in a Sakura VIP instrument. A Leica microtome was used to collect multiple 10 µm-thick sections per sample, which were mounted on positively charged glass slides and stained with hematoxylin and eosin (H & E). Images of H & E-stained sections were captured using a 20X objective on a Nikon Ti2 microscope on a Zeiss Axioscope equipped with a color camera.

### Adipose tissue analysis

Sections analyzed from the same adipose tissue block were at least 20-40 µm apart in depth, and multiple regions were imaged per section. Images were analyzed utilizing the Adiposoft plugin on ImageJ with additional manual curation to identify cell boundaries (Tholen et al., 2022). The 2D cell area was calculated as a proxy for the 3D adipocyte size. Cells located along the edges of the images or with fragmented boundaries were omitted from the analysis.

### Body Composition

Fat and lean mass distributions were measured using nucleic magnetic resonance (NMR) coupled with magnetic resonance imaging (MRI) in an EchoMRI 3-in-1 Body Composition Analyzer (EchoMRI LLC, Houston, TX). Individual mice were placed in an EchoMRI animal holding tube, allowed to acclimate to the tube for two minutes and then gently immobilized using the supplied plunger for the duration of the 90 second scan. The fat mass measured by the EchoMRI 3-in-1 Body Composition Analyzer is the mass of all the fat molecules in the body expressed as equivalent weight of canola oil. The lean mass measured is a muscle tissue mass equivalent of all the body parts containing water, excluding fat, bone minerals, and such substances which do not contribute to the NMR signal, such as hair, claws, etc.

### Hyperinsulinemic-euglycemic clamp

All procedures required for the hyperinsulinemic–euglycemic clamp were approved by the Vanderbilt University Animal Care and Use Committee. Five to seven days before the study, catheters were implanted into a jugular vein and a carotid artery of the mice for infusions and sampling, respectively, as described by Berglund et al.(Berglund et al., 2008). Insulin clamps were performed on mice fasted for 5 h using a modification of the method described by Ayala et al.(Ayala et al., 2006). After 3h of fasting, an arterial blood sample was obtained to determine natural isotopic enrichment of plasma glucose. Ninety minutes prior to initiation of the clamp, a quantitative stable isotope delivery to increase glucose isotopic enrichment above natural isotopic labelling was initiated. [6,6-^2^H_2_]glucose was primed (16 mg) and continuously infused for a 90 min equilibration and basal sampling periods (0.8 mg/kg/min in saline). The insulin clamp was initiated at t=0 min with a continuous insulin infusion (2.5-10 mU/kg body weight/min), which continued for 155 min.

Arterial glucose was clamped using a variable infusion rate of glucose + [6,6-^2^H_2_]glucose (0.08 MPE), which was adjusted based on the measurement of blood glucose at 10 min intervals during the 155 min clamp period. By mixing the glucose tracer with the unlabeled glucose infused during a clamp, deviations in arterial glucose enrichment are minimized and steady state conditions are achieved. The calculation of glucose kinetics is therefore more robust(Finegood et al., 1987). Mice received heparinized saline-washed erythrocytes from donors at a rate of 5 μl/min to prevent a fall in hematocrit. Baseline blood or plasma variables were calculated as the mean of values obtained in blood samples collected at −15 and −5 min. Blood was taken from 80–120 min for the determination of plasma enrichment. Clamp insulin was determined at *t*=100 and 120 min. At 120 min, 13µCi of 2[^14^C]deoxyglucose ([^14^C]2DG) was administered as an intravenous bolus. Blood samples were taken between 122-155min in order to determine levels of [^14^C]2DG. After the last sample, mice were anesthetized, and tissues were freeze-clamped for further analysis.

Plasma insulin was determined by RIA. Plasma C-peptide levels were determined by Luminex assay. Plasma glucose enrichments ([6,6-^2^H_2_]glucose) were assessed by GC-MS as described previously(Steele et al., 1956). Radioactivity of [^14^C]2DG in plasma samples, and [^14^C]2DG-6-phosphate in tissue samples were determined by liquid scintillation counting. Glucose appearance (Ra) and disappearance (Rd) rates were determined using steady-state equations(Steele et al., 1956). Endogenous glucose appearance (EndoRa) was determined by subtracting the GIR from total Ra. The glucose metabolic index (Rg) was calculated as previously described(Kraegen et al., 1985).

To accurately compare the relative differences in glucose uptake between muscle and fat in both groups, we first had to establish a common baseline due to the significant differences in absolute uptake between tissues. For example, visceral fat in control mice exhibited approximately 10 nmol/min/100g tissue uptake, while gastrocnemius muscle showed approximately 25 nmol/min/100g. Thus, to normalize the data and be able to compare the differences in glucose uptake between tissues, we first calculated the mean uptake for each tissue in the control mice. Thereafter, we divided the glucose uptake measured from each GC-flattened mouse by the mean glucose uptake in the corresponding tissue from control mice.

Mice were fasted for 5h before measuring plasma non-esterified fatty acid (NEFA) levels. Plasma NEFA levels were determined by the ACS-ACOD method. The percent suppression of plasma NEFA levels in Figure 4l was calculated for each mouse as (Basal - Clamp)/(Basal) *100, where “Basal” represents plasma NEFA levels measured 10 min prior to the clamp and “Clamp” represents plasma NEFA levels measured 145 minutes after the clamp was initiated.

### Western Blot Analysis

Lysates were made from flash-frozen liver tissue samples. 10 mg of frozen liver tissue was homogenized in 500 uL of RIPA buffer supplemented with 10 uL Halt protease and phosphatase inhibitor cocktail (Thermo Scientific, 78444). For further lysis and protein extraction, homogenates were incubated on ice for 30 min with intermittent vortexing and centrifuged twice thereafter at 20000g for 15 min, with each spin followed by careful collection of the intermediate clear liquid layer to avoid inclusion of tissue debris at the bottom or fatty layer at the top. The lysate collected at the end of the second spin was aliquoted to avoid repeated freeze-thaw cycles and used as needed for protein estimation by a standard BCA assay and immunoblotting. Samples were run on a 10% SDS-Polyacrylamide gel and transferred onto a 0.45 um PVDF membranes for further immunoblotting steps. Post transfer, membrane was blocked with 10% non-fat milk (dissolved in Tris buffered saline containing 0.1% Tween-20 (TBST)) for 30 min and incubated thereafter with primary antibodies diluted in TBST containing 5% BSA either for 2h at room temperature (for GAPDH) or overnight. Subsequently, membranes were washed thrice with TBST and incubated with fluorescent secondary antibodies for 75 min at room temperature, washed thrice with TBST and imaged on a LI-COR Odyssey imaging system. Following primary and secondary antibodies were used at the indicated dilutions: anti phospho-Akt Ser473 (Cell Signaling, 9271 – 1:1000), anti Akt (Cell Signaling, 9272 – 1:1000), anti - GAPDH (Cell Signaling, 5174 – 1:1000), Goat anti-rabbit Alexa Fluor 680 (Thermo Scientific, A21109 – 1:5000) and IRDye CW Goat anti-mouse 800CW (LICORbio, 926-32210 – 1:5000). Blot images were analyzed using LI-COR’s Image Studio lite software.

### Quantification and Statistical Analysis

All data are represented as mean ± SEM and representative of 3 biological replicates. Data were analyzed by unpaired t test, one-way analysis of variance (ANOVA), or two-way repeated measures ANOVA followed by Tukey’s multiple comparisons test using GraphPad Prism software. “n” indicates the number of animals per group or number of independent experiments. Results were considered significant if p < 0.05.

## Acknowledgments

This work was supported by NIH RO1-DK131432-01A1 (M.N.T.), startup funds from the Drukier Institute and Weill Cornell Medical School (M.N.T.), and NIH T32DK131957 fellowship (A.A.). We thank Tobias Meyer, Marcus Goncalves, Laura Alonso, Shannon Reilly, all members of the Teruel lab and Meyer lab (Weill Cornell Medicine), and Fredric Kraemer (Stanford) for helpful discussions. Hyperinsulinemic-euglycemic clamps were performed by the Vanderbilt Mouse Metabolic Phenotyping Center (DK020593, DK135073). The Vanderbilt Hormone Assay and Analytical Core performed insulin, c-peptide and NEFA analyses (DK020593).

## Author Contributions

A.A. and M.N.T. designed the study; A.A., S.S., A.N., J.W., E.K., L.L., A.U. and R.P. performed experiments; A.A., S.S., A.N., J.W., E.K., JL.L., R.P., and M.N.T. collected and analyzed data; T.M. gave technical support and conceptual advice. M.N.T. provided supervision and funding acquisition. A.A., S.S., and M.N.T. wrote the manuscript with input from all authors.

---

**Figure S1.**
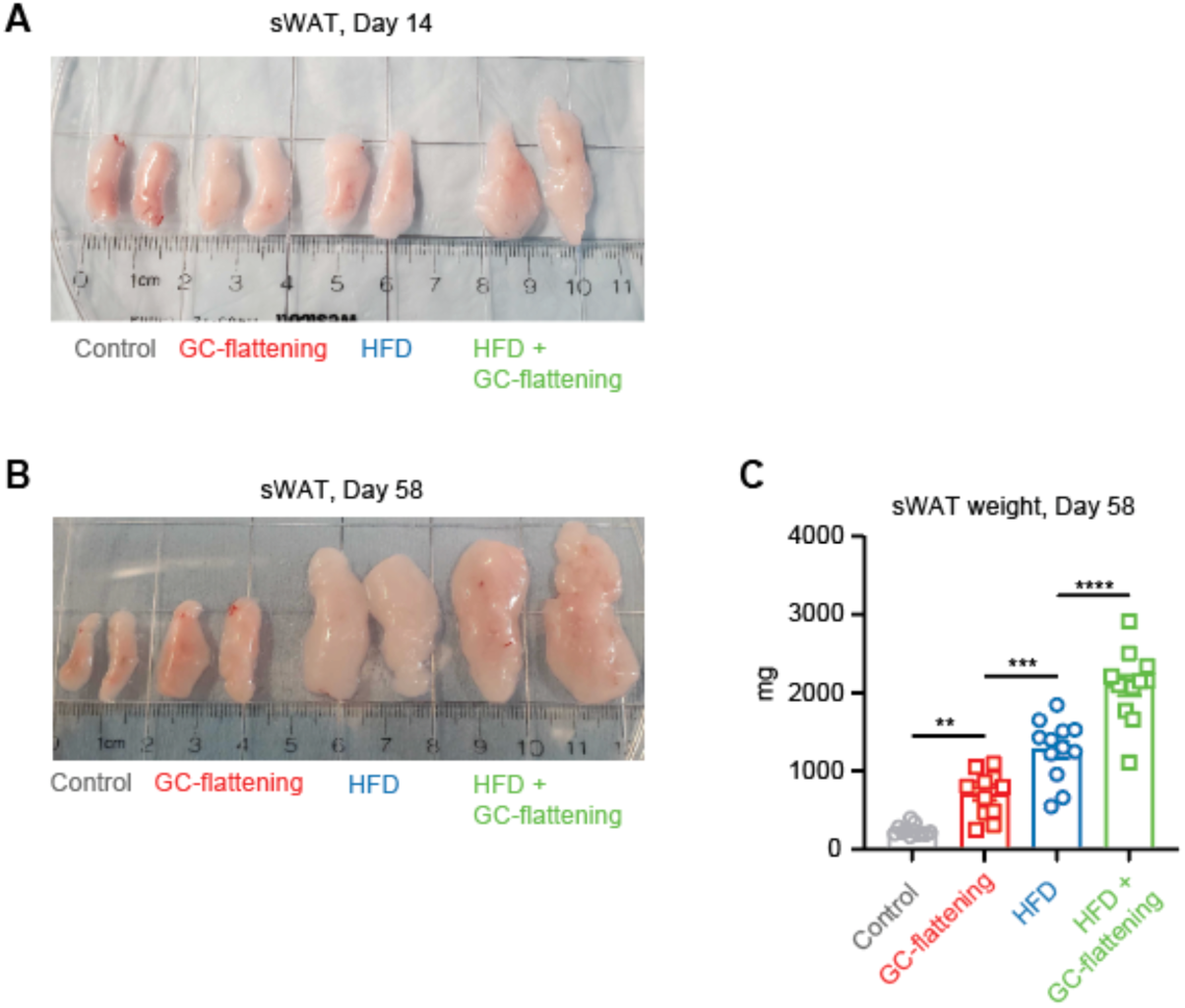
Effects of flattening of GC rhythms and high-fat diet (HFD) feeding in causing sWAT and vWAT mass increases, related to Figure 1. **(A,B)** Representative images of sWAT fat pad depots after 14 days and 58 days of treatment. **(C)** sWAT fat pad weights after 60 days of treatment Data show means ± SEM, n=12 mice per group pooled from two experimental repeats. **(C)** One-way ANOVA followed by Tukey’s post-hoc test for multiple comparisons. **p < 0.01, ***p < 0.001, ****p < 0.0001.

**Figure S2.**
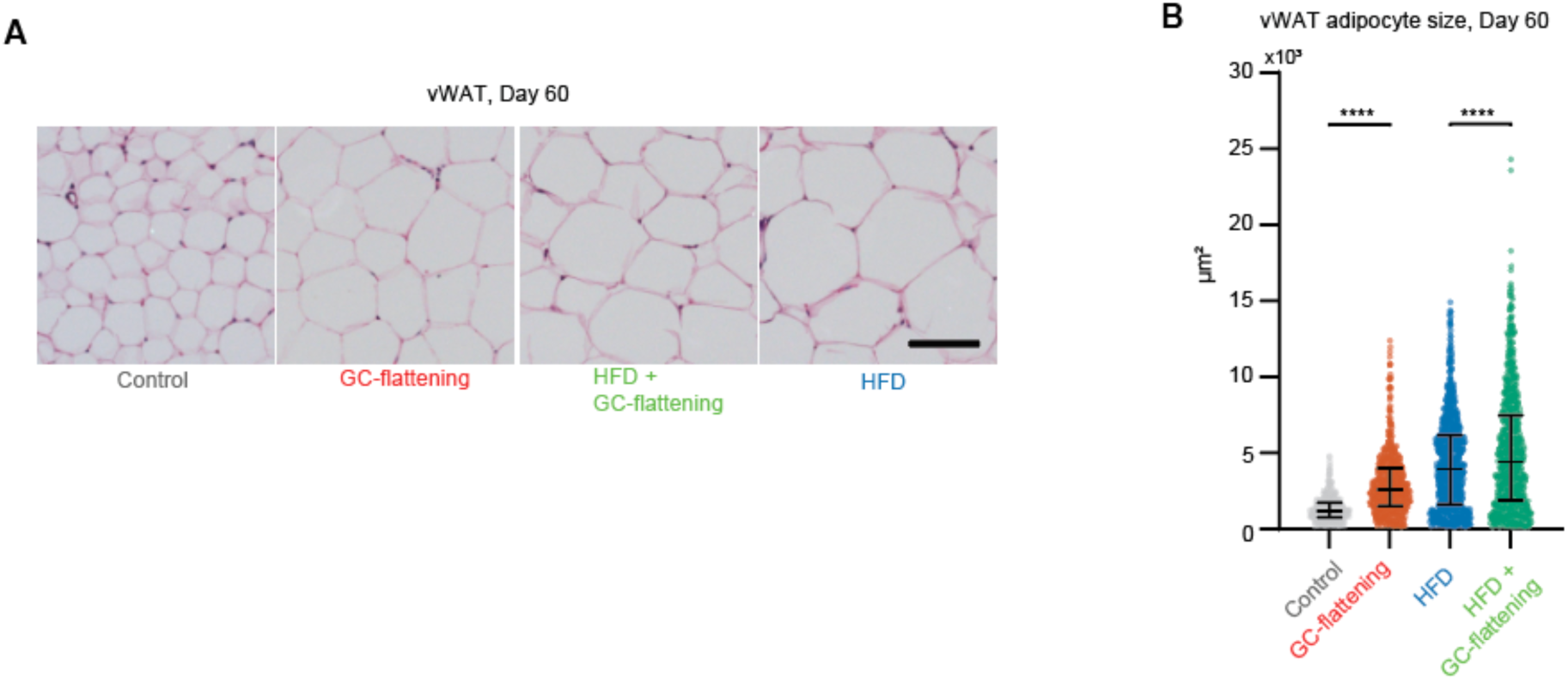
Flattening of GC rhythms and high-fat diet (HFD) feeding have additive effects in causing vWAT adipocyte hypertrophy, related to Figure 1. **(A),** Representative H&E images of vWAT depots following 8 weeks of treatment. Scale bar = 100 µm. **(B)** Scatter plot showings the size distribution of individual adipocytes with median and 25th and 75^th^ percentiles marked. Data was collected from 6 mice per group. 700-1000 total number of cells analyzed from 6 mice per group. **(B)** one-way ANOVA followed by Tukey’s post-hoc test for multiple comparisons****p < 0.0001.

**Figure S3.**
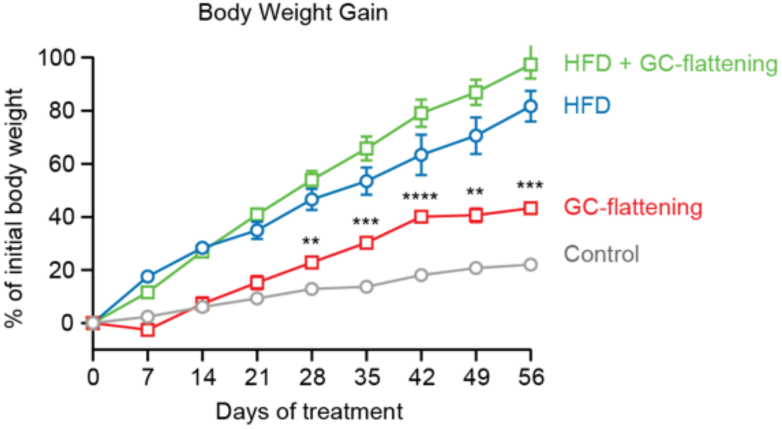
Flattening of GC rhythms in mice promotes body weight gain on both chow and high-fat diet, related to Figure 2. Body weights of control or GC-flattened mice fed a chow or HFD were measured throughout the 8 weeks of treatments, and the differences in body weights for each animal at every time point were calculated with respect to its initial body weight and plotted. Data show means ± SEM, n=5 mice per group.

**Figure S4.**
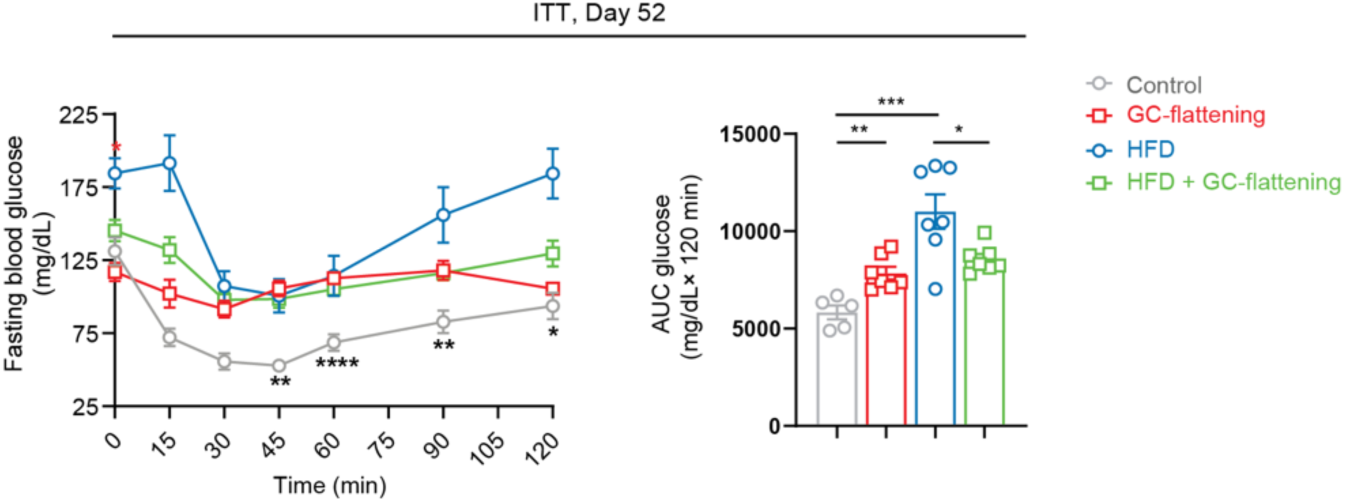
GC-flattening causes insulin resistance independent of diet, related to Figure 4. An intraperitoneal insulin tolerance test (ITT) was performed following 52 days of treatment. Left panel shows blood glucose levels measured in each treatment group during the ITT. The panel on the right shows the cumulative area under the curve (AUC) during the ITT. Data show means ± SEM, n=5-7 mice per group. Unpaired t-test was used to compare for statistical differences between indicated treatment groups. *p < 0.05, **p < 0.01, ***p < 0.001.

**Figure S5.**
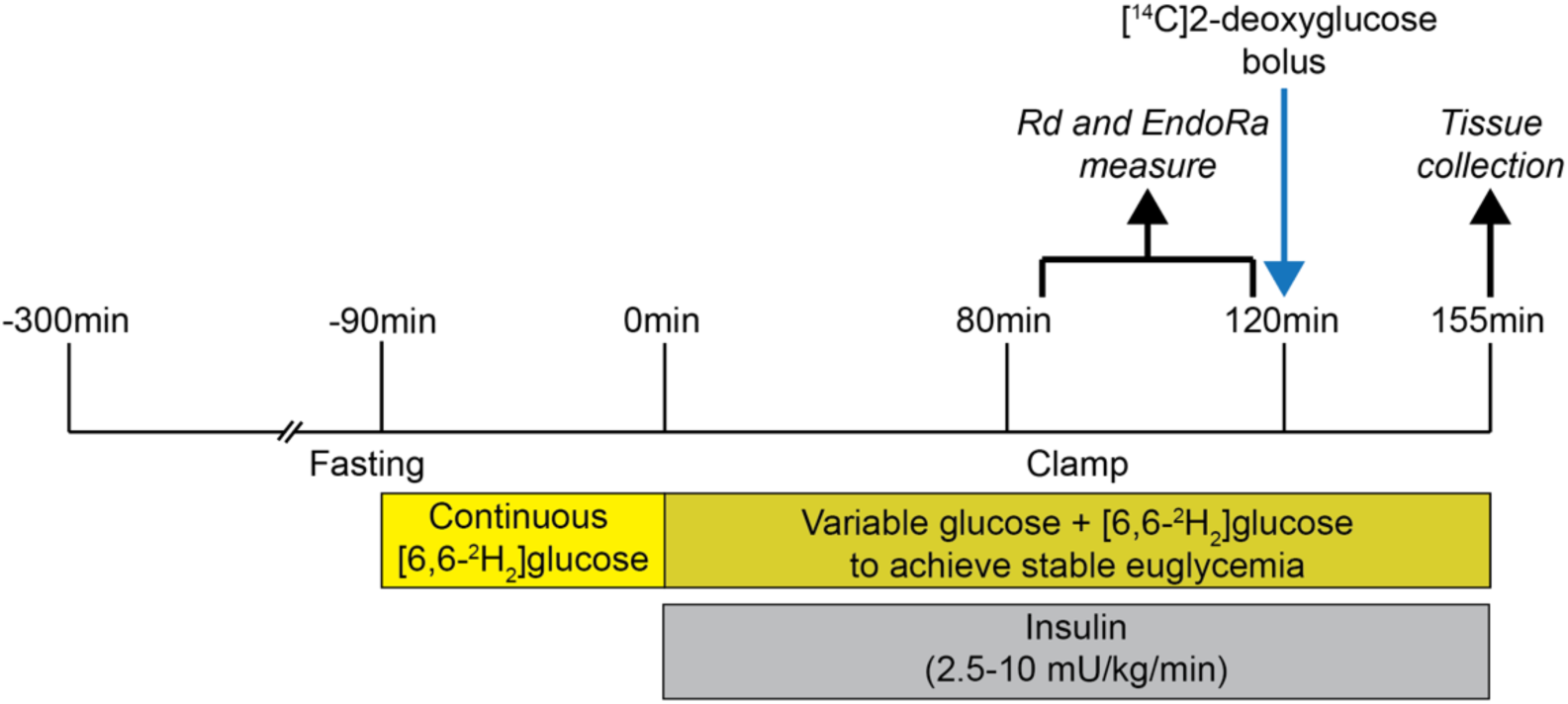
Timeline of the hyperinsulinemic-euglycemic clamp studies, related to Figure 4.

**Figure S6.**
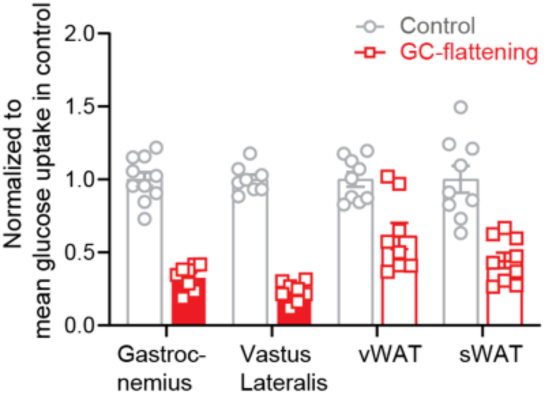
Hyperinsulinemic-euglycemic clamp experiments show GC-flattening reduces glucose uptake into tissues, related to Figure 6. For tissue specific glucose uptake measurements mice received an intravenous bolus of radioactive 2[^14^C]deoxyglucose at 120 min and sacrificed for tissue dissections 35 min later. Glucose uptake into muscle (gastrocnemius and vastus lateralis) and fat (vWAT and sWAT) tissues of GC-flattened mice normalized to the mean glucose uptake in corresponding tissues of control mice.

## REFERENCES

Almeida, D.M., J. Rush, J. Mogle, J.R. Piazza, E. Cerino, and S.T. Charles. 2023. Longitudinal change in daily stress across 20 years of adulthood: Results from the national study of daily experiences. Dev Psychol. 59:515–523. doi:10.1037/dev0001469.

Alquier, T., and V. Poitout. 2018. Considerations and guidelines for mouse metabolic phenotyping in diabetes research. Diabetologia. 61:526–538. doi:10.1007/s00125-017-4495-9.

Ayala, J.E., D.P. Bracy, O.P. McGuinness, and D.H. Wasserman. 2006. Considerations in the Design of Hyperinsulinemic-Euglycemic Clamps in the Conscious Mouse. Diabetes. 55:390–397. doi:10.2337/diabetes.55.02.06.db05-0686.

Bahrami-Nejad, Z., M.L. Zhao, S. Tholen, D. Hunerdosse, K.E. Tkach, S. van Schie, M. Chung, and M.N. Teruel. 2018. A Transcriptional Circuit Filters Oscillating Circadian Hormonal Inputs to Regulate Fat Cell Differentiation. Cell Metabolism. 27:854–868.e8. doi:10.1016/j.cmet.2018.03.012.

Berglund, E.D., C.Y. Li, G. Poffenberger, J.E. Ayala, P.T. Fueger, S.E. Willis, M.M. Jewell, A.C. Powers, and D.H. Wasserman. 2008. Glucose Metabolism In Vivo in Four Commonly Used Inbred Mouse Strains. Diabetes. 57:1790–1799. doi:10.2337/db07-1615.

Blüher, M. 2019. Obesity: global epidemiology and pathogenesis. Nat Rev Endocrinol. 15:288–298. doi:10.1038/s41574-019-0176-8.

Broussard, J.L., and E. Van Cauter. 2016. Disturbances of sleep and circadian rhythms: novel risk factors for obesity. Current Opinion in Endocrinology & Diabetes and Obesity. 23:353–359. doi:10.1097/MED.0000000000000276.

Campbell, J.E., A.J. Peckett, A.M. D’souza, T.J. Hawke, and M.C. Riddell. 2011. Adipogenic and lipolytic effects of chronic glucocorticoid exposure. American Journal of Physiology-Cell Physiology. 300:C198–C209. doi:10.1152/ajpcell.00045.2010.

Chaput, J.-P., A.W. McHill, R.C. Cox, J.L. Broussard, C. Dutil, B.G.G. Da Costa, H. Sampasa-Kanyinga, and K.P. Wright. 2023. The role of insufficient sleep and circadian misalignment in obesity. Nat Rev Endocrinol. 19:82–97. doi:10.1038/s41574-022-00747-7.

Corkey, B.E. 2012. Banting Lecture 2011: Hyperinsulinemia: Cause or Consequence? Diabetes. 61:4–13. doi:10.2337/db11-1483.

Dallman, M., S. Akana, S. Bhatnagar, M. Bell, and A. Strack. 2000. Bottomed out: metabolic significance of the circadian trough in glucocorticoid concentrations. International Journal of Obesity. 24:S40–S46.

Finegood, D.T., R.N. Bergman, and M. Vranic. 1987. Estimation of Endogenous Glucose Production During Hyperinsulinemic-Euglycemic Glucose Clamps. 36.

Gasparini, S.J., M.-C. Weber, H. Henneicke, S. Kim, H. Zhou, and M.J. Seibel. 2016. Continuous corticosterone delivery via the drinking water or pellet implantation: A comparative study in mice. Steroids. 116:76–82. doi:10.1016/j.steroids.2016.10.008.

Geißler, C., C. Krause, A.-M. Neumann, J.H. Britsemmer, N. Taege, M. Grohs, M. Kaehler, I. Cascorbi, A.G. Lewis, R.J. Seeley, H. Oster, and H. Kirchner. 2022. Dietary induction of obesity and insulin resistance is associated with changes in Fgf21 DNA methylation in liver of mice. The Journal of Nutritional Biochemistry. 100:108907. doi:10.1016/j.jnutbio.2021.108907.

Harno, E., C. Sefton, J.R. Wray, T.-J. Allen, A. Davies, A.P. Coll, and A. White. 2021. Chronic glucocorticoid treatment induces hepatic lipid accumulation and hyperinsulinaemia in part through actions on AgRP neurons. Sci Rep. 11:13776. doi:10.1038/s41598-021-93378-3.

Hodes, G.E., B.R. Brookshire, T.E. Hill-Smith, S.L. Teegarden, O. Berton, and I. Lucki. 2012. Strain differences in the effects of chronic corticosterone exposure in the hippocampus. Neuroscience. 222:269–280. doi:10.1016/j.neuroscience.2012.06.017.

Hodson, L., and P.J. Gunn. 2019. The regulation of hepatic fatty acid synthesis and partitioning: the effect of nutritional state. Nat Rev Endocrinol. 15:689–700. doi:10.1038/s41574-019-0256-9.

Imamura, F., R. Micha, S. Khatibzadeh, S. Fahimi, P. Shi, J. Powles, and D. Mozaffarian. 2015. Dietary quality among men and women in 187 countries in 1990 and 2010: a systematic assessment. The Lancet Global Health. 3:e132–e142. doi:10.1016/S2214-109X(14)70381-X.

Janssen, J.A.M.J.L. 2021. Hyperinsulinemia and Its Pivotal Role in Aging, Obesity, Type 2 Diabetes, Cardiovascular Disease and Cancer. IJMS. 22:7797. doi:10.3390/ijms22157797.

Joseph, J.J., and S.H. Golden. 2017. Cortisol dysregulation: the bidirectional link between stress, depression, and type 2 diabetes mellitus: Role of cortisol in stress, depression, and diabetes. Ann. N.Y. Acad. Sci. 1391:20–34. doi:10.1111/nyas.13217.

Karatsoreos, I.N., S.M. Bhagat, N.P. Bowles, Z.M. Weil, D.W. Pfaff, and B.S. McEwen. 2010. Endocrine and Physiological Changes in Response to Chronic Corticosterone: A Potential Model of the Metabolic Syndrome in Mouse. Endocrinology. 151:2117–2127. doi:10.1210/en.2009-1436.

Koch, C.E., B. Leinweber, B.C. Drengberg, C. Blaum, and H. Oster. 2017. Interaction between circadian rhythms and stress. Neurobiology of Stress. 6:57–67. doi:10.1016/j.ynstr.2016.09.001.

Kraegen, E.W., D.E. James, A.B. Jenkins, and D.J. Chisholm. 1985. Dose-response curves for in vivo insulin sensitivity in individual tissues in rats. American Journal of Physiology-Endocrinology and Metabolism. 248:E353–E362. doi:10.1152/ajpendo.1985.248.3.E353.

Kroon, J., M. Schilperoort, W. In Het Panhuis, R. Van Den Berg, L. Van Doeselaar, C.R.C. Verzijl, N. Van Trigt, I.M. Mol, H.H.C.M. Sips, J.K. Van Den Heuvel, L.L. Koorneef, R.J. Van Der Sluis, A. Fenzl, F.W. Kiefer, S. Vettorazzi, J.P. Tuckermann, N.R. Biermasz, O.C. Meijer, P.C.N. Rensen, and S. Kooijman. 2021. A physiological glucocorticoid rhythm is an important regulator of brown adipose tissue function. Molecular Metabolism. 47:101179. doi:10.1016/j.molmet.2021.101179.

Lang, P., S. Hasselwander, H. Li, and N. Xia. 2019. Effects of different diets used in diet-induced obesity models on insulin resistance and vascular dysfunction in C57BL/6 mice. Sci Rep. 9:19556. doi:10.1038/s41598-019-55987-x.

Lian, C.-Y., Z.-Z. Zhai, Z.-F. Li, and L. Wang. 2020. High fat diet-triggered non-alcoholic fatty liver disease: A review of proposed mechanisms. Chemico-Biological Interactions. 330:109199. doi:10.1016/j.cbi.2020.109199.

Luijten, I.H.N., K. Brooks, N. Boulet, I.G. Shabalina, A. Jaiprakash, B. Carlsson, A.W. Fischer, B. Cannon, and J. Nedergaard. 2019. Glucocorticoid-Induced Obesity Develops Independently of UCP1. Cell Reports. 27:1686–1698.e5. doi:10.1016/j.celrep.2019.04.041.

Meijer, O.C., S. Kooijman, J. Kroon, and E.M. Winter. 2023. The importance of the circadian trough in glucocorticoid signaling: a variation on B-flat. Stress. 26:2275210. doi:10.1080/10253890.2023.2275210.

Micha, R., S. Khatibzadeh, P. Shi, S. Fahimi, S. Lim, K.G. Andrews, R.E. Engell, J. Powles, M. Ezzati, D. Mozaffarian, and on behalf of the Global Burden of Diseases Nutrition and Chronic Diseases Expert Group (NutriCoDE). 2014. Global, regional, and national consumption levels of dietary fats and oils in 1990 and 2010: a systematic analysis including 266 country-specific nutrition surveys. BMJ. 348:g2272–g2272. doi:10.1136/bmj.g2272.

Miller, G.E., E. Chen, and E.S. Zhou. 2007. If it goes up, must it come down? Chronic stress and the hypothalamic-pituitary-adrenocortical axis in humans. Psychological Bulletin. 133:25–45. doi:10.1037/0033-2909.133.1.25.

Monteiro, C.A., J. -C. Moubarac, G. Cannon, S.W. Ng, and B. Popkin. 2013. Ultra-processed products are becoming dominant in the global food system. Obesity Reviews. 14:21–28. doi:10.1111/obr.12107.

Odeleye, O.E., M. de Courten, D.J. Pettitt, and E. Ravussin. 1997. Fasting Hyperinsulinemia Is a Predictor of Increased Body Weight Gain and Obesity in Pirna Indian Children. 46:5.

Peckett, A.J., D.C. Wright, and M.C. Riddell. 2011. The effects of glucocorticoids on adipose tissue lipid metabolism. Metabolism. 60:1500–1510. doi:10.1016/j.metabol.2011.06.012.

Popkin, B.M., L.S. Adair, and S.W. Ng. 2012. Global nutrition transition and the pandemic of obesity in developing countries. Nutrition Reviews. 70:3–21. doi:10.1111/j.1753-4887.2011.00456.x.

Sapolsky, R.M. 2021. Glucocorticoids, the evolution of the stress-response, and the primate predicament. Neurobiology of Stress. 14:100320. doi:10.1016/j.ynstr.2021.100320.

Sapolsky, R.M., L.C. Krey, and B.S. McEWEN. 1986. The Neuroendocrinology of Stress and Aging: The Glucocorticoid Cascade Hypothesis. 7:18.

Schweiger, M., M. Romauch, R. Schreiber, G.F. Grabner, S. Hütter, P. Kotzbeck, P. Benedikt, T.O. Eichmann, S. Yamada, O. Knittelfelder, C. Diwoky, C. Doler, N. Mayer, W. De Cecco, R. Breinbauer, R. Zimmermann, and R. Zechner. 2017. Pharmacological inhibition of adipose triglyceride lipase corrects high-fat diet-induced insulin resistance and hepatosteatosis in mice. Nat Commun. 8:14859. doi:10.1038/ncomms14859.

Spiegel, K., R. Leproult, and E. Van Cauter. 1999. Impact of sleep debt on metabolic and endocrine function. The Lancet. 354:1435–1439. doi:10.1016/S0140-6736(99)01376-8.

Steele, R., J.S. Wall, R.C. De Bodo, and N. Altszuler. 1956. Measurement of Size and Turnover Rate of Body Glucose Pool by the Isotope Dilution Method. American Journal of Physiology-Legacy Content. 187:15–24. doi:10.1152/ajplegacy.1956.187.1.15.

Steensels, S., J. Qiao, Y. Zhang, K.M. Maner-Smith, N. Kika, C.D. Holman, K.E. Corey, W.C. Bracken, E.A. Ortlund, and B.A. Ersoy. 2020. Acyl-Coenzyme A Thioesterase 9 Traffics Mitochondrial Short-Chain Fatty Acids Toward De Novo Lipogenesis and Glucose Production in the Liver. Hepatology. 72:857–872. doi:10.1002/hep.31409.

Surwit, R.S., C.M. Kuhn, C. Cochrane, J.A. McCUBBIN, and M.N. Feinglos. 1988. Diet-Induced Type II Diabetes in C57BL/6J Mice. 37.

Taylor, S.R., S. Ramsamooj, R.J. Liang, A. Katti, R. Pozovskiy, N. Vasan, S.-K. Hwang, N. Nahiyaan, N.J. Francoeur, E.M. Schatoff, J.L. Johnson, M.A. Shah, A.J. Dannenberg, R.P. Sebra, L.E. Dow, L.C. Cantley, K.Y. Rhee, and M.D. Goncalves. 2021. Dietary fructose improves intestinal cell survival and nutrient absorption. Nature. 597:263–267. doi:10.1038/s41586-021-03827-2.

Tholen, S., R. Patel, A. Agas, K.M. Kovary, A. Rabiee, H.T. Nicholls, E. Bielczyk-Maczyńska, W. Yang, F.B. Kraemer, and M.N. Teruel. 2022. Flattening of circadian glucocorticoid oscillations drives acute hyperinsulinemia and adipocyte hypertrophy. Cell Reports. 39:111018. doi:10.1016/j.celrep.2022.111018.

Turner, N., G.M. Kowalski, S.J. Leslie, S. Risis, C. Yang, R.S. Lee-Young, J.R. Babb, P.J. Meikle, G.I. Lancaster, D.C. Henstridge, P.J. White, E.W. Kraegen, A. Marette, G.J. Cooney, M.A. Febbraio, and C.R. Bruce. 2013. Distinct patterns of tissue-specific lipid accumulation during the induction of insulin resistance in mice by high-fat feeding. Diabetologia. 56:1638–1648. doi:10.1007/s00125-013-2913-1.

Vegiopoulos, A., and S. Herzig. 2007. Glucocorticoids, metabolism and metabolic diseases. Molecular and Cellular Endocrinology. 275:43–61. doi:10.1016/j.mce.2007.05.015.

Weitzman, E.D., D. Fukushima, C. Nogeire, H. Roffwarg, T.F. Gallagher, and L. Hellman. 1971. Twenty-four Hour Pattern of the Episodic Secretion of Cortisol in Normal Subjects. The Journal of Clinical Endocrinology & Metabolism. 33:14–22. doi:10.1210/jcem-33-1-14.

